# Escape Steering by Cholecystokinin Peptidergic Signaling

**DOI:** 10.1101/2021.12.08.471862

**Authors:** Lili Chen, Yuting Liu, Pan Su, Wesley Hung, Haiwen Li, Ya Wang, Zhongpu Yue, Minghai Ge, Zhengxing Wu, Yan Zhang, Peng Fei, Li-Ming Chen, Louis Tao, Heng Mao, Mei Zhen, Shangbang Gao

## Abstract

Escape is an evolutionarily conserved and essential avoidance response. Considered to be innate, most studies on escape responses focused on hard-wired circuits. We report here that peptidergic signaling is an integral and necessary component of the *Caenorhabditis elegans* escape circuit. Combining genetic screening, electrophysiology and calcium imaging, we reveal that a neuropeptide NLP-18 and its cholecystokinin receptor CKR-1 enable the escape circuit to execute a full omega (Ω) turn, the last motor step where the animal robustly steers away from its original trajectory. We demonstrate *in vivo* and *in vitro* that CKR-1 is a Gα_q_ protein coupled receptor for NLP-18. *in vivo*, NLP-18 is mainly secreted by the gustatory sensory neuron (ASI) to activate CKR-1 in the head motor neuron (SMD) and the turn-initiating interneuron (AIB). Removal of NLP-18, removal of CKR-1, or specific knockdown of CKR-1 in SMD or AIB neurons lead to shallower turns hence less robust escape steering. Consistently, elevation of head motor neuron (SMD)’s Ca^2+^ transients during escape steering is attenuated upon the removal of NLP-18 or CKR-1. *in vitro*, synthetic NLP-18 directly evokes CKR-1-dependent currents in oocytes and CKR-1-dependent Ca^2+^ transients in SMD. Thus, cholecystokinin signaling modulates an escape circuit to generate robust escape steering.

## Introduction

Neuromodulators, including a rich variety of neuropeptides, reconfigurate neural circuits by transforming the intrinsic property of neurons and altering synaptic strength between neuronal wiring (Grillner and Jessell, 2009; Marder et al., 2014). Small invertebrate circuits have provided fundamental insights on conserved circuit principles and neuromodulation that underlie locomotory behaviors (Friedrich, 2013; Friesen and Kristan, 2007; Katz, 2016). Escape, a nociceptive response to steer away from a threat that an animal encounters during foraging, has been a particularly attractive model of ethologically important sensorimotor transformations. Escape occurs fast and exhibits robust stereotype across species (Chalfie and Jorgensen, 1998; Harris-Warrick, 2011; Wang et al., 2020). For example, looming stimuli, which represent objects on a collision course, initiates robust escape responses across vertebrates and invertebrates (Fotowat and Gabbiani, 2011).

*Caenorhabditis elegans* (*C. elegans*) employs a dedicated circuit for escape, from predacious fungi (Maguire et al., 2011; Pirri and Alkema, 2012) to nociceptive stimuli (Chalfie et al., 1985; Hilliard et al., 2005). Its escape steering consists of a highly orchestrated sequence of motor actions (Pirri et al., 2009; Wang et al., 2020). A strong mechanical touch to its head, for example, initiates a reversal followed by a head-led, ventral-biased turn that allows the animal to reorient its foraging trajectory. During the turn, the nose of the animal touches the middle ventral body and glides to the tail, resulting a turn that resembles an omega (termed Ω turn). The escape motor sequence finishes with this head-led forward movment (**Figure 1A**). Ω turn allows robust steering away fromthe stimuli.

**Figure 1.**
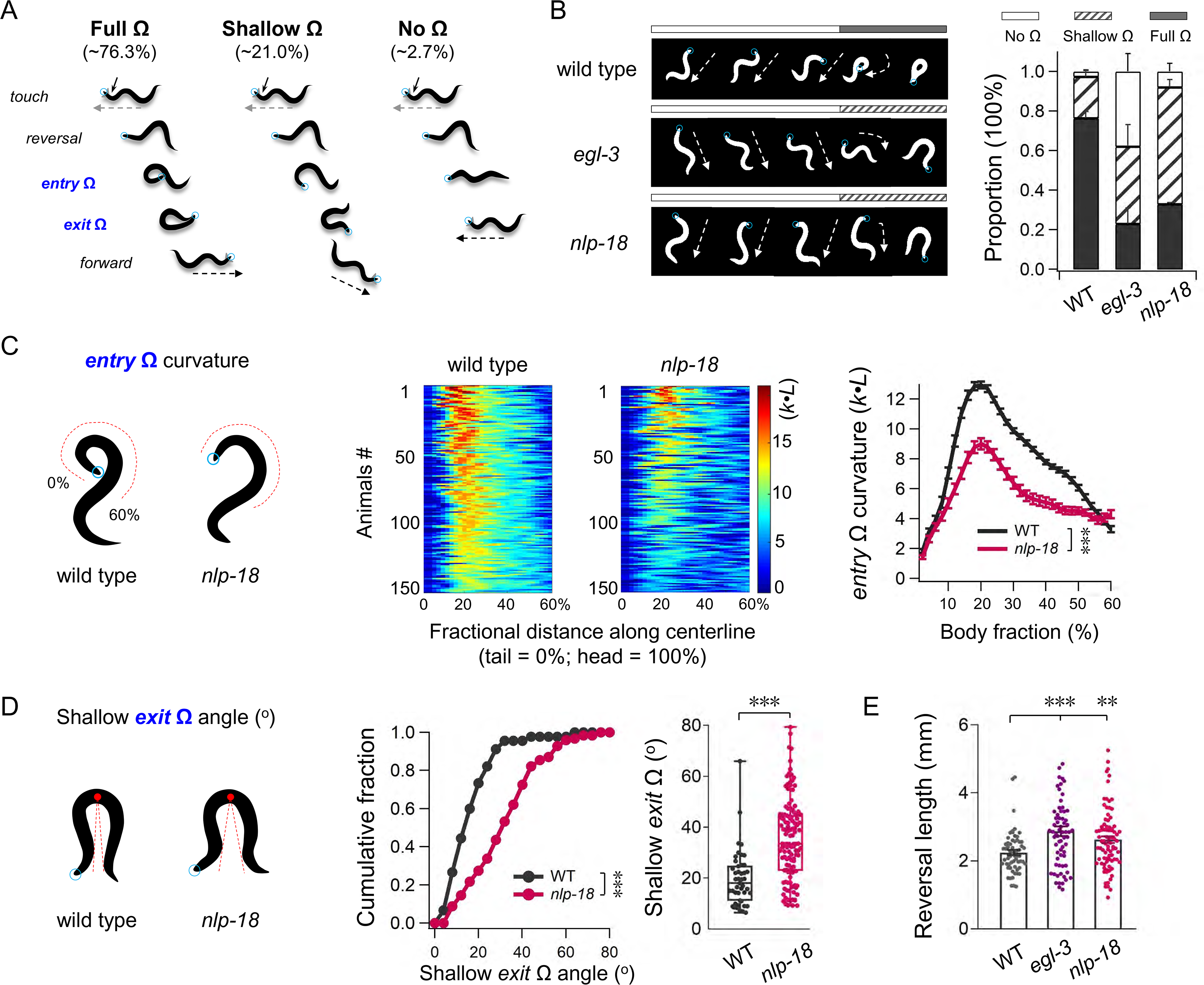
Mutant in Neuropeptide like Peptide Gene nlp-18 disrupted the Robustness of Omega Turn. (**A**) A sketch diagram of escape response by head touch, including reversal, omega (Ω) turn and following forward movement. The action of Ω turn is accompanied by two steps: *entry* Ω and *exit* Ω. Blue circles denote the head, dash arrows denote the forward orientation. (**B**) Sequential snapshots of representative Ω body postures and quantification of the full Ω proportion in different genotypes. Mutants in *egl-3* and *nlp-18* display obvious Ω turn defects, exhibiting shallower Ω were mostly observed (n ≥ 5 trials, at least 30 animals each trial). Statistical analysis of proportion was performed with the Fisher’s exact test. (**C**) *Left*, schematic representation of the *entry* Ω curvature. *Right*, representative color maps and average curvatures of the *entry* Ω in wild type and *nlp-18* mutant animals (n ≥ 150 animals). \*\*\**P* < 0.001, Two-way ANOVA test. (**D**) *Left*, schematic representation of the shallow *exit* Ω angle. The *exit* Ω angle was defined as the angle from the deepest point in the bend to the closest points anterior and posterior of the animal. *Right*, the distribution and quantification of all shallow *exit* Ω angle in wild type and *nlp-18* mutant animals, respectively (n ≥ 45 animals). \*\*\**P* < 0.001, Kolmogorov-Smirnov test (*Left*) and Mann-Whitney test (*Right*). (**E**) Harsh touch induced reversal length in different genotypes (n ≥ 60 animals). Each point represents the angle or reversal length from one animal. \*\**P* < 0.01, \*\*\**P* < 0.001, Mann-Whitney test. All data are expressed as mean ± SEM.

Detailed wiring diagram (White et al., 1986) guided the laser ablation studies that delineate neuronal components of the *C. elegans* navigation motor circuit (Chalfie et al., 1985; Gray et al., 2005; Wicks et al., 1996). Key circuit components for the escape response are mechano- and chemo-sensory neurons that detect noxious stimuli, descending synapses that activate head motor neurons for neck muscles, and ventral-cord-projecting premotor interneurons that regulate motor neurons along the body. Separate motor modules, consisted of interneurons and motor neurons, execute the sequential escape motor steps: reversal (interneurons AVA and AIB), turn (motor neurons SMD and RIV), and forward (interneurons RIB and AVB). Chemical and electrical synaptic connections between neurons of these modules generate feedforward excitation and mutual inhibition, generating a set of motor sequence with flexibility in their transitions (Croll, 1975; Kawano et al., 2011; Piggott et al., 2011; Pirri and Alkema, 2012; Pirri et al., 2009; Wakabayashi et al., 2004; Wang et al., 2020).

Regulation of the turning amplitude and frequency of the Ω turn determine effectiveness of escape. This involves neurons of and outside of core escape circuit. With reduced food signals, a gustatory sensory neuron (ASI) and an interneuron (AIY) reduce the frequency of the Ω turn (Gray et al., 2005). Activation of AIB during long reversals increases turning frequency (Gordus et al., 2015; Wang et al., 2020). The amplitude of Ω turn is coded by the excitatory head motor neurons (SMD), and the ventral turning bias of Ω turn is specified by the motor neurons for ventral neck muscles (RIV) (Gray et al., 2005). Some Ω turn regulation is context-specific. Strong activation of an interneuron (RIM) inhibits head movements (Alkema et al., 2005; Pirri and Alkema, 2012; Pirri et al., 2009) during long reversals, but facilitates body bending (Donnelly et al., 2013; Kagawa-Nagamura et al., 2018) after the animal initiates the Ω turn. Essential neuronal signaling for escape motor sequences has mostly focused on chemical and electrical signaling. These pioneering works significantly promote the escape ‘hard-wired’ circuit.

Neuromodulation modifies properties and states of neurons and their connections. Modulators include a small number of monoamines and a large repertoire of peptides. Many monoamines exert conserved and global modulation, such as dopamine for the reward circuit (Schultz, 2002). In contrast, neuropeptides exhibit vast varieties, with astonishing functional specificity for neurons and circuit configurations (Grillner and Jessell, 2009; Marder and Bucher, 2001). An exemplary example is the crustacean stomatogastric ganglion, where over 50 neuropeptides act on a small set of neurons and synapses to produce different rhythmic output patterns (Blitz and Nusbaum, 2008; Ye et al., 2013).

*C. elegans* genome may encode up to 250 neuropeptides, which consist of three main classes: the FMRF-amide-related (FLP), insulin-like (INS), and non-insulin/non-FMRF-amide-related but neuropeptide-like protein (NLP) (Husson and Schoofs, 2007; Li and Kim, 2010; Pierce et al., 2001). Neuropeptides activate G-protein coupled receptors (GPCRs) to initiate intracellular signaling cascades that exert long-lasting and long-range effects (Nassel, 2009). In *C. elegans*, more than 125 potential neuropeptide GPCRs were predicted (Frooninckx et al., 2012; Janssen et al., 2010). Several neuropeptides have been found to affect locomotion (Bhardwaj et al., 2018; Hu et al., 2011; Janssen et al., 2008; Lim et al., 2016; Meelkop et al., 2012; Oranth et al., 2018). However, neuropeptides or receptors that are directly involved in execution or regulation of escape have not been reported.

Combining behavioral studies, Ca^2+^ imaging, heterogenous reconstitution, and *in situ* neuronal activity recordings, we identified a peptide NLP-18 and its receptor CKR-1, a G α_q_-protein coupled receptor, form an essential signaling pathway for Ω turn. We show that through NLP-CKR-1 signaling pathway, the gustatory neuron (ASI) activates the interneuron (AIB)-to-motor neuron (SMD) subcircuit to allow Ω turn. Like synaptic transmission, this neuromodulatory signaling plays a selective and necessary role in an innate and robust motor action.

## Results

### Escape steering requires peptidergic signaling

To examine the circuit underpinning escape steering, we first adapted an escape assay (Li et al., 2011) that robustly induces stereotype responses to afford easy quantification of turning. Using a platinum wire to deliver a single head touch per adult animal cultured on a thin layer of food (Methods), 97% wild type (N2) animals evoked a three-step motor response: reversal, turn, and forward with a different heading angle (**Figure 1A, B, and Supplementary Movie S1**). Among them, 76.3±3.3% exhibited robust escape steering, defined by connecting reversal and forward movement with a head reorientation called a full Ω turn: the head bends, touches and glides off the posterior half of the ventral body. 21.0±3.4% exhibited less robust head reorientation, connecting reversal and forward with a shallow Ω turn, where the head bends towards but not touching the body (**Figure 1A, D**). A very small fraction of animals (2.7±0.4%) exhibited no turn (no Ω), and even rarely (2 in total 189 assays) they did a δ-shaped turn where the head bends, touches the anterior body, and crosses over to exit at the dorsal body (Broekmans et al., 2016). Propensity and properties of turns are presented by three parameters: fraction of full versus shallow Ω turns, peak curvature of anterior body during turning (*entry* curvature), and angle of deepest middle body bending at the end of turning (*exit* angle) (Methods). This assay led to reliable responses (≥ 3 trials, at least 30 animals each trial).

To assess a potential involvement of peptidergic signaling, we compared escape steering of wild type animals to mutant animals that cannot synthesize active neuropeptides. Active neuropeptides are derived from precursor proteins processed by a pro-protein convertase (PC) EGL-3 and a carboxypeptidase E (CPE) (EGL-21) (Jacob and Kaplan, 2003; Kass et al., 2001). In the absence of either processing enzyme, propensity of full Ω turns was reduced, to 23.0±8% (*egl-3*, *P* < 0.001 against N2) and 16.7±3% (*egl-21, P <* 0.001 against N2), respectively (**Figure 1B, Supplementary Figure S1A**). CAPS/UNC-31 is required for dense-core vesicle fusion hence neuropeptide release. In the absence of CAPS, propensity of full Ω turn was similarly decreased (*unc-31* 27.7±4.9%*, P <* 0.001 against N2). During escape, propensity of full Ω turns is positively correlated to the length of reversal (Zhao et al., 2003). Reduced proportion Ω turn in peptidergic signaling mutants may be consequential to a reduced reversal length. However, our assay induced a slight increased reversal length between wild type animals and animals that cannot synthesize active neuropeptide (*egl-3* 2.87±0.14 mm, N2 2.25±0.08 mm; *P* < 0.001) (**Figure 1E**).

*C. elegans* genome encodes three peptide families: NLPs, FLPs and INS (Li and Kim, 2008), most of which require PC/EGL-3 processing. Insulin-like peptides function through an INS-family receptor (DAF-2) and a FOXO transcription factor (DAF-16). Reducing insulin-like signaling, by either functional reduction of DAF-2 or elimination of DAF-16 did not alter propensity of full Ω turn during escape steering (**Supplementary Figure S1A**).

Collectively, these results implicate a critical and specific requirement of non-ILP neuropeptides signaling underlie robust escape steering.

### NLP-18 is critical for robust escape steering

We screened loss-of-function mutants for 20 FLP and 12 NLP encoding genes using our assay. Four exhibited statistically significant reduction of escape responses with full Ω turns. Among them, removal of an NLP encoding gene, *nlp-18*, led to the most consistent and closest degree of reduction (*nlp-18* 32.8±1.4%, *P* < 0.01 against N2) to the effect of removing EGL-3 (**Supplementary Figure S1B**). Removal of three FLP-encoding genes also reduced the propensity of full Ω turns, but their effect was either modest (*flp-18* and *flp-20*), or inconsistent between trials (*flp-1*) (**Supplementary Figure S1C**).

The *nlp-18* gene (**Figure 2A**) encodes a propeptide, which is processed into five mature peptides called NLP-18a-e, respectively (**Figure 2B**). *nlp-18(ok1557)* is a null allele that removes the partial propeptide coding sequence (**Figure 2A**). *nlp-18* null mutants entered the turn with significantly reduced curvature than wild type animals (**Figure 1C**). Not only had a much higher proportion of *nlp-18* mutant animals exited with a shallow Ω turn (∼ 60%) than wild type animals (∼ 20%) (**Figure 1B)**, shallow Ω turns occurred at a higher exit angle for *nlp-18* mutants than wild type animals (**Figure 1D, Supplementary Movie S2**). Similar to *egl-3* mutants, duration of reversal in *nlp-18* mutants was slightly increased during evoked escape (2.62±0.09 mm vs 2.25±0.08 mm for N2 wildtype, *P* < 0.01) (**Figure 1E**). Spontaneous velocity and propensity of directional movement were unchanged in *nlp-18* mutants (**Supplementary Figure S3A, B**). These results demonstrate that *nlp-18* may specifically promote a full Ω turn.

**Figure 2.**
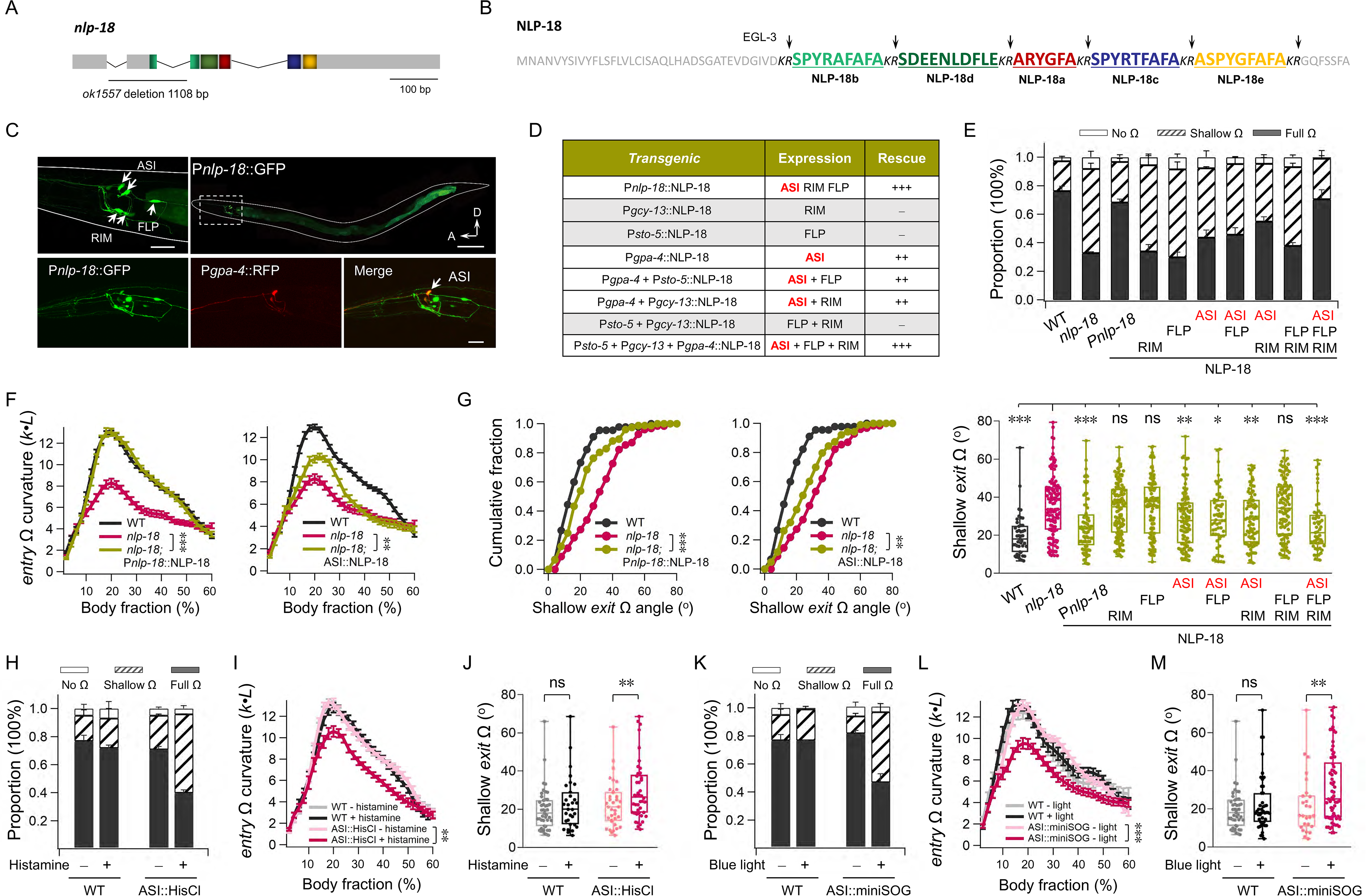
ASI neurons are critical for nlp-18 modulated robust escape steering. (**A**) The gene structure of *nlp-18*(*ok1557*) with 1108 bp deletion indication. (**B**) *nlp-18* encodes a neuropeptide precursor that harbors five putative neuropeptides, which are flanked by the dibasic *KR* (arrow) cleaving sites. Colors label the predicted neuropeptide sequences (red NLP-18a, green NLP-18b, blue NLP-18c, dark green NLP-18d, golden NLP-18e). (**C**) Expression pattern of *nlp-18*. *Up*, endogenous *nlp-18* promoter driven GFP was observed in three head neurons (ASI, FLP and RIM) and intestine. Scale bar, left 20 μm, right 100 μm. *Bottom*, the co-localization of P*nlp-18*::GFP and P*gpa-4*::RFP in ASI neurons. Scale bar, 20 μm. Worm orientation, A anterior, D dorsal. (**D**) Summary the rescue of Ω turn deficiency by the expression of NLP-18 in different neurons. ASI are the critical functional neurons for NLP-18. _ no rescue, + slight rescue, ++ moderate rescue, +++ full rescue. (**E**) Propensity of the head touch induced escape responses (no Ω, shallow Ω and full Ω) from wild type, *nlp-18* and the rescue strains. Full Ω proportion could be properly restored by expression of NLP-18 in ASI (n ≥ 150 animals). Statistical analysis of proportion was performed with the Fisher’s exact test. (**F, G**) The *entry* Ω curvature and shallow *exit* Ω angle were restored by self-promoter driven NLP-18 and ASI specific expression of NLP-18 (n ≥ 45 animals). Two-way ANOVA tested in F. Kolmogorov-Smirnov test was used in G (*Left*). \*\**P* < 0.01 and \*\*\**P* < 0.001. (**H**-**J**) Silence of ASI neurons by histamine (10 mM) significantly reduced the full Ω proportion (H) and the *entry* Ω curvature (I), increased the shallow *exit* Ω angle (J). Fisher’s exact test in H. Two-way ANOVA tested in I, \*\**P* < 0.01. Mann-Whitney test in J. ns, not significant, \*\**P* < 0.01. (**K**-**M**) Ablation of ASI neurons significantly reduced the full Ω proportion (K) and the *entry* Ω curvature (L), increased the shallow *exit* Ω angle (M). Fisher’s exact test in K. Two-way ANOVA tested in L, \*\*\**P* < 0.001. Mann-Whitney test in M. ns, not significant, \*\**P* < 0.01. (H-M) n ≥ 3 trials, each trial with at least 30 worms were tested. Data are presented as mean ± SEM.

Requirement of *nlp-18* for robust escape steering is not unique to mechanical stimuli. When we evoked escape by either an aversive metal (**Supplementary Figure S1D**) or high osmolarity (**Supplementary Figure S1E**), removal of *nlp-18* led to reduced propensity for full Ω turn and increased propensity for shallow Ω turn, as well as shallow Ω turns with reduced entry curvature and increased exit angles. Thus, *nlp-18* is an inherent component of the escape circuit to promote deep turning.

### NLP-18 promotes escape steering mainly from the ASI sensory neuron

To pinpoint NLP-18’s role in the escape circuit, we first assessed its cellular origin. A transcriptional reporter for *nlp-18* (Methods) revealed strong expression in the intestine and three pairs of neurons: a gustatory sensory neuron (ASI), a mechanical sensory neuron (FLP), and an interneuron (RIM) (**Figure 2C, Supplementary Figure S2B**).

When NLP-18 expression was restored by this promoter, *nlp-18* mutants’ turning defects were fully rescued (**Figure 2D, Supplementary Figure S2C, D**). This result implicates functional sufficiency of *nlp-18* among these cells.

We then systematically tested the functional requirement for individual cells using exogenous promoters. Driving NLP-18 expression in the intestine did not change escape behaviors of *nlp-18* mutants (not shown). Expression of NLP-18 in the mechanosensory neuron (FLP) or interneuron (RIM) did not rescue *nlp-18* mutants either. By contrast, driving NLP-18 expression in the gustatory sensory neuron (ASI) significantly rescued *nlp-18* mutant animal’s turning defects of in our assay, with an increased propensity of full Ω turns (**Figure 2D, E**), increased entry body curvature of the turn (**Figure 2F**), and decreased exit angles at the end of shallow Ω turns (**Figure 2G**). Restoring NLP-18 in three neurons together led to an effect qualitatively similar to that of ASI-specific NLP-18 expression, with modest but statistically significant improvement (**Figure 2E-G**).

Implication of this result - NLP-18 promotes robust escape steering from the ASI neurons - surprised us. A previous laser ablation study found that ASI inhibit the frequency of short reversals and Ω turns in a food-signal dependent manner (Gray et al., 2005). To clarify the role of ASI during escape steering, we examined the effect of silencing ASI neurons using histamine-gated chloride channel (HisCl) (Pokala et al., 2014). Exposure of animals that ectopically and specifically expressed HisCl in ASI to histamine led to reduced full Ω turns and increased shallow Ω turns (**Figure 2H**), as well as reduced entry curvature and larger exit angle for shallow Ω turns (**Figure 2I, G**). We observed similar effects (**Figure 2K-M**) when we ablated ASI using mini singlet oxygen generator (miniSOG) (Qi et al., 2012; Shu et al., 2011). *nlp-18* mutant’s escape steering defect was also not affected by the absence of food (**Supplementary Figure S3E-G**). These results confirm that the gustatory sensory neurons ASI are required for NLP-18-dependent robust escape steering.

### A cholecystokinin receptor CKR-1 promotes robust escape steering

Molecular mapping of receptors is necessary to delineate peptidergic signaling. To identify NLP-18’s physiological receptors, we started by examining deletion mutants for GPCRs using our assay. ∼150 predicted *C. elegans* GPCRs have significant sequence homologies to peptidergic receptors in other systems (Frooninckx et al., 2012). Among them, we screened 31 with significant homology to mammalian peptidergic receptors, including the CKR (cholecystokinin receptor), NPR (neuropeptide receptor) and NMUR (neuromedin U receptor) families, as well as receptors predicted to bind NLPs (Frooninckx et al., 2012).

Among them, removal of the *ckr-1* gene led to severely reduced escape steering **(Supplementary Figure S3C**). CKR-1 is closely related in sequence to the cholecystokinin receptor CCKR-1/CCKAR (**Figure 3A, B, Supplementary Figure S3D**), receptors implicated satiety, anxiety, and gall bladder contraction (Berna et al., 2007). *ckr-1* mutant animals’ escape defects are highly reminiscent to *nlp-18* mutants, with reduced full Ω turn propensity (**Figure 3C**), increased entry curvature (**Figure 3D**) and reduced exit angle (**Figure 3E**). Compared to *nlp-18* mutants, the severity of *ckr-1*’s defect was slightly reduced. Reversal length during evoked escape was not altered in *ckr-1* mutants (2.22±0.09 mm vs N2 2.25±0.08 mm, *P* > 0.05) (**Figure 3F**). Spontaneous velocity and partition between directional movements were also unchanged in *ckr-1*, *nlp-18* or *nlp-18; ckr-1* mutants (**Supplementary Figure S3A, B**). Similar to *nlp-18* mutants, *ckr-1*’s steering defect is not dependent on food signals (**Supplementary Figure S3E-G**).

**Figure 3.**
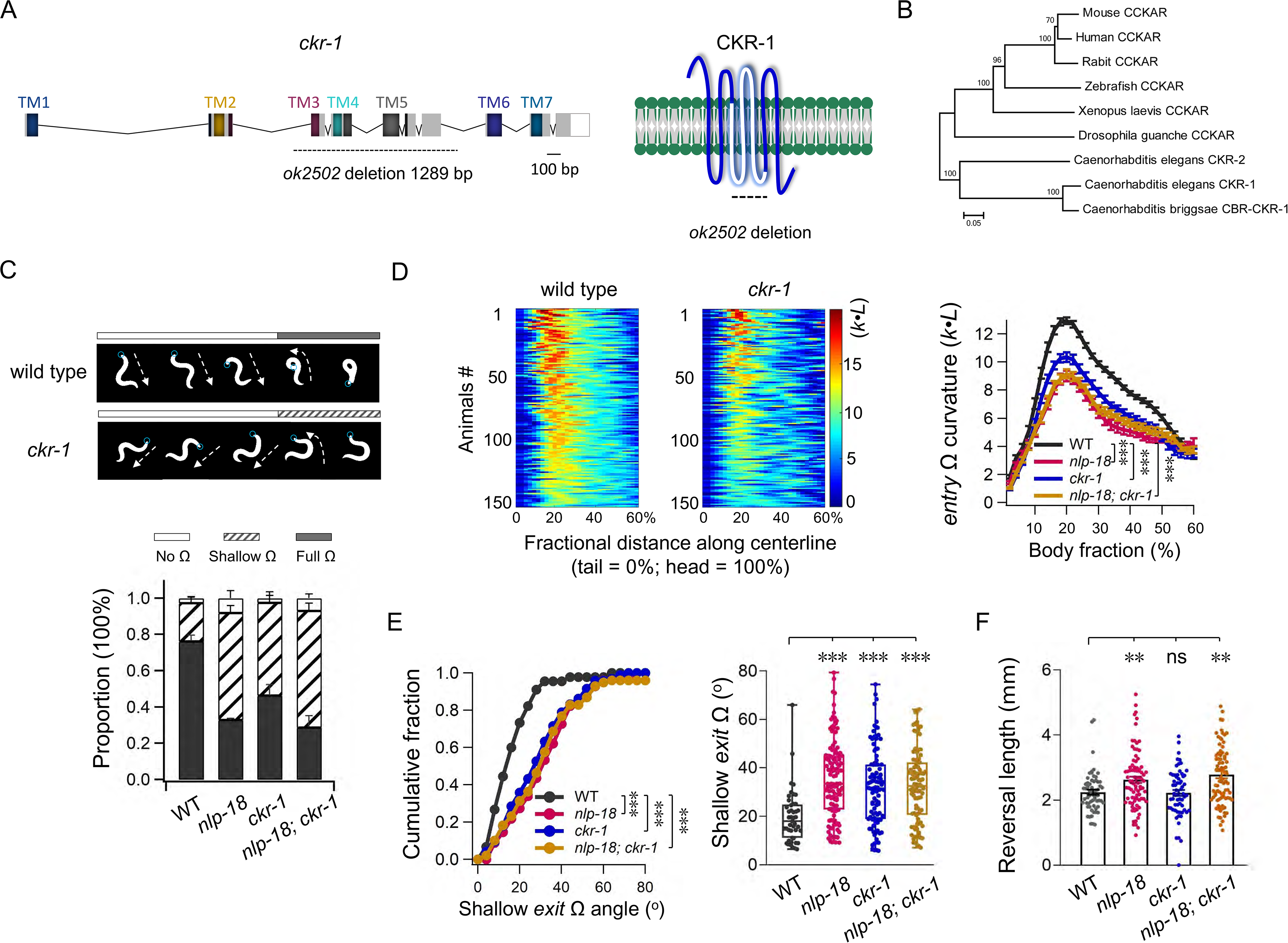
CKR-1, a cholecystokinin receptor, is required for robust escape steering. (**A**) The gene and protein structure of *ckr-1(ok2502)* with 1289 bp deletion indication. Seven predicted transmembrane (TM) domains were marked. (**B**) Phylogenetic tree (analyzed by MEGA 6.60) demonstrates that *C. elegans ckr-1* encodes a cholecystokinin A receptor. (**C**) *Up*, sequential snapshots of representative body postures of the Ω turn in wild type and *ckr-1*. Blue points denote the head. *Bottom*, propensity of head touch evoked escape responses (no Ω, shallow Ω and full Ω) in wild type, *nlp-18*, *ckr-1* and *nlp-18; ckr-1* double mutants (≥ 5 trials, at least 30 animals each trial). Fisher’s exact test in C (*Bottom*). (**D**) Representative color maps and quantification of the *entry* Ω curvature in wild type, *ckr-1* and *nlp-18; ckr-1* mutants (n ≥ 150 animals). \*\*\**P* <0.001, Two-way ANOVA test. (**E**) *ckr-1* and *nlp-18; ckr-1* show similar increased shallow *exit* Ω angle phenotype of *nlp-18* (n ≥ 45 animals). Kolmogorov-Smirnov test in E *(Left).* Mann-Whitney test was used in E *(Right).* \*\*\**P* <0.001. (**F**) Harsh touch induced reversal length in different genotypes (n ≥ 60 animals). Mann-Whitney test, ns, not significant, \*\**P* < 0.01. Data are expressed as mean ± SEM.

These results indicate that CKR-1 and NLP-18 function in the same signaling pathway to promote robust escape steering.

### ckr-1 promotes steering from the motor (SMD) and inter (AIB) neurons of the escape circuit

To map where NLP-18-CKR-1 signaling pathway may take place, we first sought where CKR-1 resides in the escape circuit. A functional transcriptional reporter for CKR-1 revealed strong expression in many neurons and weak expression in the intestine (**Figure 4A**). CKR-1-expressing neurons include head motor neurons SMD and RME (**Figure 4B**), interneurons AIB and RIM, peptidergic neurons RIS and body motor neurons A, B, and D. All are involved in motor behaviors. Among them, SMD, RME, AIB and RIM have been implicated in motor sequences specific for escape (Alkema et al., 2005; Gray et al., 2005; Hendricks et al., 2012; Kaplan et al., 2020; Pirri et al., 2009; Shen et al., 2016; Wang et al., 2020).

**Figure 4.**
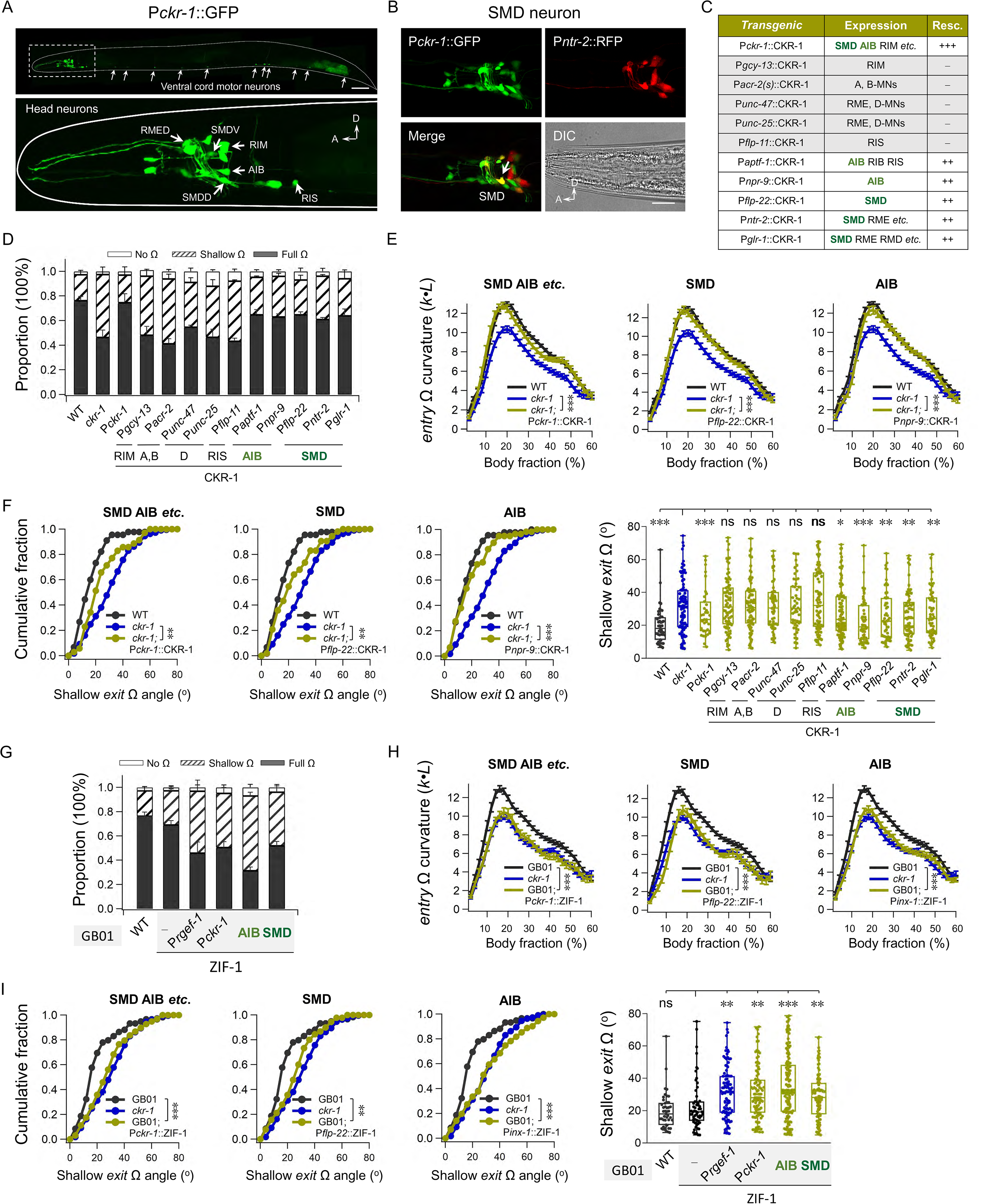
CKR-1 regulates escape steering robustness from SMD/AIB neurons. (**A**) Expression pattern of *ckr-1*. *Up, ckr-1* expresses in the *C. elegans* nervous system including the head and ventral cord neurons, scale bar 100 μm. *Bottom*, representative head neurons (SMD, AIB, RME, RIM, RIS and other neurons, scale bar 20 μm). A anterior, D dorsal. (**B**) The P*ckr-1*::GFP expression in SMD neurons. (**C**) Summary of the rescue potency of Ω turn deficiency that expression of CKR-1 in different neurons, from which SMD and AIB exhibit critical functions of *ckr-1*. _ no rescue, + slight rescue, ++ moderate rescue, +++ full rescue. (**D**) Distribution of the head touch induced escape responses (no Ω, shallow Ω and full Ω) in distinct transgenic strains (n ≥ 5 trials, at least 30 animals each trial). The proportion of full Ω was properly restored by the expression of CKR-1 in SMD and AIB neurons, but not in other neurons. Statistical analysis of proportion was performed with the Fisher’s exact test. (**E, F**) The *entry* Ω curvature and shallow *exit* Ω angle in *ckr-1* mutant were rescued by the expression of CKR-1 in SMD and AIB neurons. (n ≥ 45 animals). Two-way ANOVA test were used in E. Shallow exit Ω angle cumulative fraction was tested by Kolmogorov-Smirnov in F (*Left*) and Mann-Whitney test in F (*Right*). ns, not significant, \**P* < 0.05, \*\**P* < 0.01 and \*\*\**P* < 0.001. (**G**) Composition of the head touch induced escape responses (no Ω, shallow Ω and full Ω) in wild type, *ckr-1* and CKR-1 knock down strains with the expression of neuronal specific ZIF-1 in GB01 *gaaIs1* background (≥ 5 trials, at least 30 animals each trial). Statistical analysis of proportion was performed with the Fisher’s exact test. (**H, I**) The *entry* Ω curvature and shallow *exit* Ω angle in GB01 were reduced by the expression of ZIF-1 in SMD and AIB neurons. (I) Scatter diagram and quantification of the shallow *exit* Ω angles from different knock down strains (n ≥ 46 animals). Two-way ANOVA test were used in H. Shallow exit Ω angle cumulative fraction was tested by Kolmogorov-Smirnov in I (*Left*), and Mann-Whitney in I (*Right*). ns, not significant, \*\**P* < 0.01 and \*\*\**P* < 0.001. Data are expressed as mean ± SEM.

We identified key neurons through which CKR-1 promotes escape steering by restoration of its expression using exogenous promoters that overlap with those cells. Driving CKR-1 expression in either SMD (by P*flp-22,* P*ntr-2* or P*glr-1*) or AIB (by P*aptf-1* or P*npr-9*) led to a robust rescue of escape steering defects in *ckr-1* mutant animals (**Figure 4C**), evaluated by the proportion of full Ω turns (**Figure 4D**), entry curvature (**Figure 4E**) and exit angle (**Figure 4F**). Driving CKR-1 expression in all other neurons or the intestine did not lead to rescue (**Figure 4C, D, F**; **Supplementary Figure S4A, B**).

Restoring CKR-1 in either SMD or AIB led to near full degree of rescue, which implicates redundancy of CKR-1-signaling in the escape circuit. Alternatively, it may be an artifact of exogenously restored expression, because exogenous promoters did not deliver CKR-1 at its physiological level, and in some experiments, introduced ectopic CKR-1 signaling in the circuit. To further verify where endogenous CKR-1 functions from, we examined the effect of depletion of CKR-1 protein from SMD or AIB, using a repurposed non-neuronal ubiquitin system (Armenti et al., 2014). Briefly, we tagged the endogenous *ckr-1* locus with ZF1, an E3-recognition target signal for the ZIF-1 ligase. When ZIF-1 is expressed in targeted neurons, endogenous CKR-1 proteins are degraded cell-specifically (Methods).

Insertion of the ZF1 motif into the *ckr-1* locus did not change behaviors (**Figure 4G**), development or physiology. When ZIF-1 was expressed in either SMD or AIB neurons in *ckr-1*::ZF1 animals (GB01), they exhibited less robust escape steering, where turning included less full Ω turns, the entry curvature when starting the turn was decreased, whereas the *exit* angle was significantly increased (**Figure 4G-I**). These animals qualitatively resembled the escape defects of *ckr-1* genetic mutant with a statistically significant small difference. Critically, results of the rescue and mimic experiments corroborate that CKR-1-mediated signaling can promote deep turning through either the AIB interneuron or the SMD head motor neuron.

### CKR-1 encodes a G _q_ protein-coupled receptor

CKR-1’s mammalian homologue CCKR-1 is a G protein-coupled receptor. *C. elegans* has homologs for four mammalian Gα subtypes, EGL-30/Gα_q_, GSA-1/Gα_s_, GOA-1/Gα_i/o_ and GPA-12/Gα_12/13_ (Frooninckx et al., 2012; Jansen et al., 1999). EGL-30 binds and activates EGL-8, the PLCβ homolog, which converts PIP_2_ to DAG and IP_3_ to initiate ER Ca^2+^ release. Among four Gα subtypes, only EGL-30/Gα_q_ is required for the full Ω proportion (**Figure 5A**). Full Ω proportion was also reduced in *egl-8* mutant, to the same level of *egl-30*. Entry Ω curvature, as well as the shallow *exit* Ω angle, were significantly reduced in *egl-8* mutant, supporting that Gα_q_ signaling is required for the escape steering. Importantly, removing CKR-1 in *egl-30* or *egl-8* mutant did not further enhance their escape defects (**Figure 5B, D, E)**, further supporting that *ckr-1* and *egl-30*, *egl-8* are in the same signaling pathway. Thus, CKR-1 functions as a Gα_q_ coupled receptor to regulate escape steering (**Figure 5C**).

**Figure 5.**
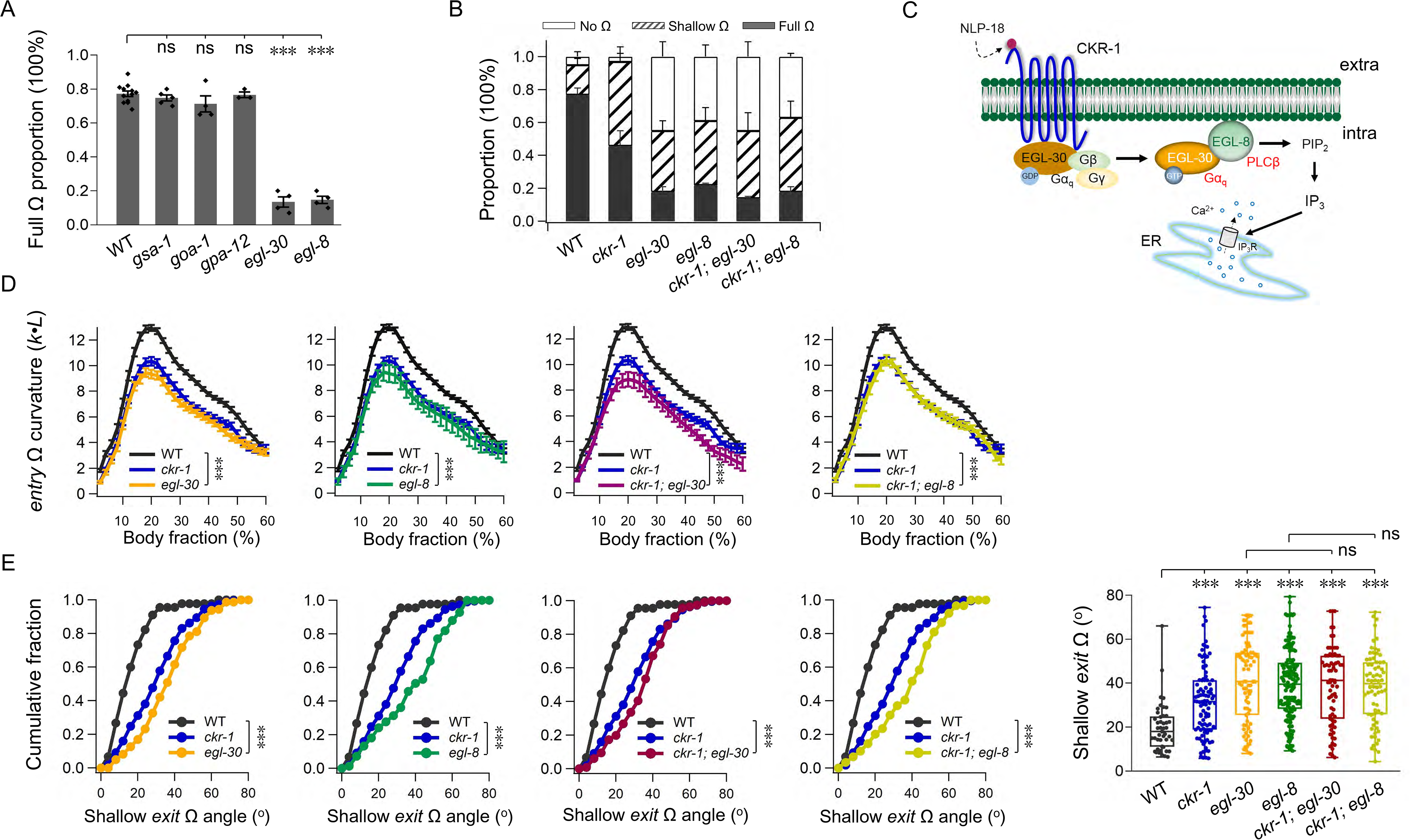
CKR-1 is a Gα_q_-protein coupled receptor. (**A**) Quantification of the full Ω proportion in wild type and various G-protein signaling mutants. The full Ω proportion was specifically reduced in *egl-30* and *egl-8* mutants. ns, not significant, \*\*\**P* < 0.001, Student’s *t*-test. (**B**) Composition of the head touch induced escape responses (no Ω, shallow Ω and full Ω) in wild type, *ckr-1* and respective mutants. *ckr-1* could not further reduce the full Ω proportion in *egl-30* and *egl-8* mutants. (A, B) n ≥ 3 trials, at least 30 animals each trial. Statistical analysis of proportion was performed with the Fisher’s exact test. (**C**) Schematics of putative CKR-1 G-protein coupling signaling pathway. CKR-1 is coupled by the Gα_q_ protein. (**D, E**) The *entry* Ω curvature and shallow *exit* Ω angle recapitulate the level of *ckr-1* mutant in *egl-30*, *egl-8* single mutant and *ckr-1; egl-30*, *ckr-1; egl-8* double mutants. Scatter diagram and quantification of the shallow *exit* Ω angle from wild type, *ckr-1* mutant and the G-protein signaling mutants. *ckr-1* could not further reduce the shallow *exit* Ω angle in *egl-30* and *egl-8* mutant backgrounds (n ≥ 46 animals). Two-way ANOVA test were used to access the statistical difference in D. Shallow exit Ω angle cumulative fraction was tested by Kolmogorov-Smirnov in E (*Left*) and Mann-Whitney in E *(Right)*. ns, not significant, \*\*\**P* < 0.001. Data are expressed as mean ± SEM.

### SMD motor neurons exhibit NLP-18-CKR-1-dependent activity increase during the Ω turn

Two neurons we found to be involved in CKR-1 signaling play different roles in *C. elegans* motor behaviors. AIB’s Ca^2+^ dynamics correlates with the transition between reversal and turning (Wang et al., 2020), whereas SMD’s periodic Ca^2+^ undulation correlated with ventral and dorsal head bends (Hendricks et al., 2012; Kaplan et al., 2020).

We measured the SMD activity in freely moving animals by a genetic calcium sensor GCaMP6s (Methods). As reported in above studies, SMDV and SMDD exhibited calcium signals that correlates with ventral and dorsal head bending during foraging and turning. Animals turn both ventrally and dorsally. Here, we focused on activity of SMDD neuron and their correlation with dorsal turns (**Figure 6A)**. When evoked to escape, SMDD exhibited periodic Ca^2+^ dynamics (peak value denoted as R Ca^2+^) (**Figure 6B**), which positively correlated with the dorsal head bending (**Figure 6B, C**). After the movement transited into a full Ω turn, we noticed a further increase of Ca^2+^ signals (peak value denoted as Ω Ca^2+^) (**Figure 6B, Supplementary movie S3**). The further increase of Ω Ca^2+^ from R Ca^2+^ was significant (**Figure 6D**) and expected: a deeper head bend during the Ω turn reflects higher SMD activity and increased excitatory inputs to head muscles.

**Figure 6.**
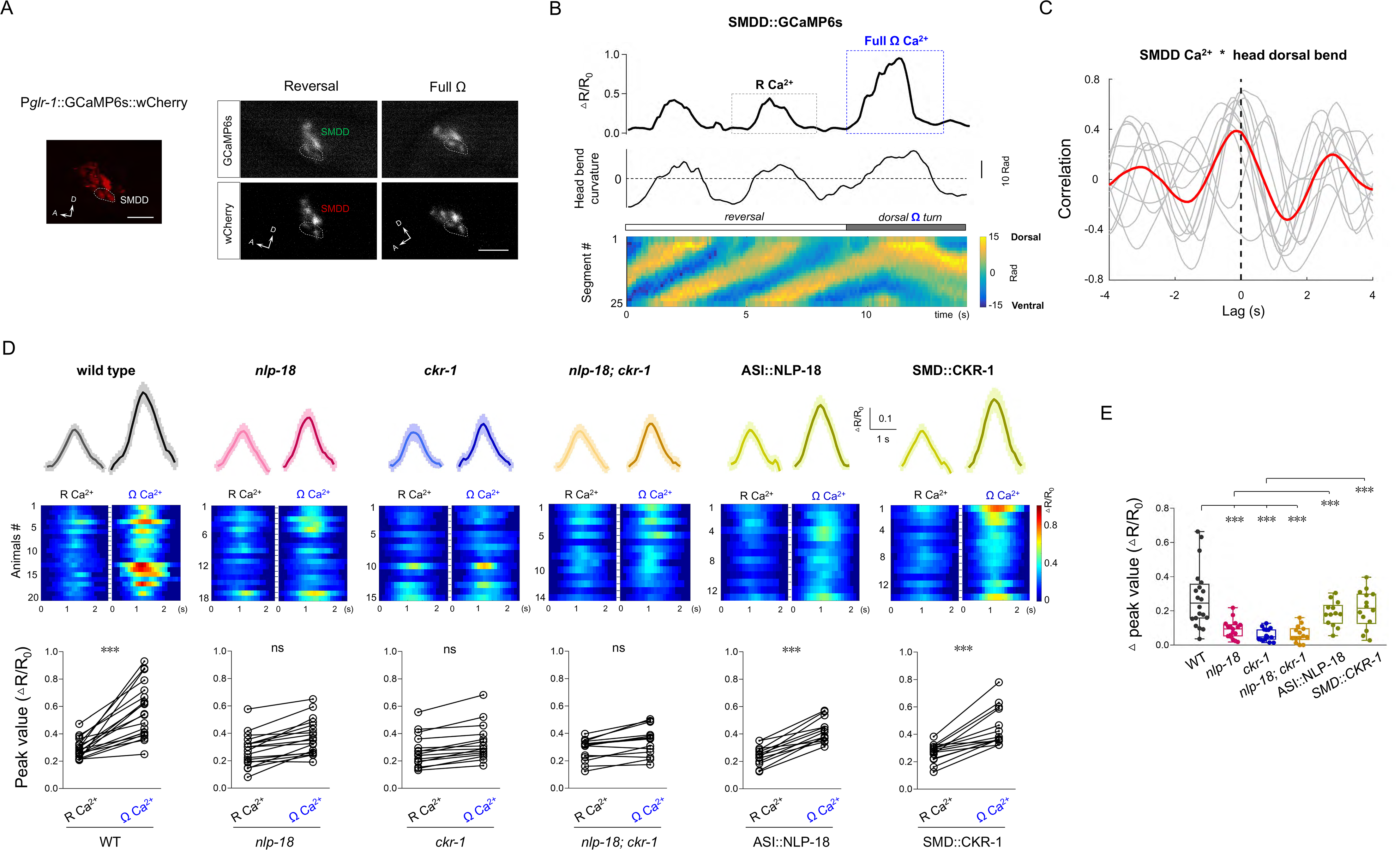
The Ω Ca^2+^ of SMD neurons was reduced in escape steering mutants. (**A**) Representative images showing fluorescence of GCaMP6s and wCherry in the SMDD soma of P*glr-1*::GCaMP6s::wCherry transgenic worms during reversal and full Ω. *Left*, a pair of SMDD::wCherry identified by 60× objective. A anterior, D dorsal. Scale bar: 10 μm. (**B)** *Up*, representative Ca^2+^ transient trace of SMDD and head bend curvature in the process of reversal and turn from a free-behaving animal. △R/R0 = normalized GCaMP/wCherry ratio. *Bottom*, posture kymograms. (**C**) Cross-correlation between SMDD Ca^2+^ and head dorsal bend. Faint lines indicate the results from individual animal and the red line indicate mean value, n = 10. (**D**) *Up*, the average traces of the last R Ca^2+^ and Ω Ca^2+^ in different genotypes. *Middle*, the representative color maps of the corresponding Ca^2+^ activities. *Bottom*, the peak Ca^2+^ values of the adjacent R Ca^2+^ and Ω Ca^2+^ in different genotypes. ns, not significant, \*\*\**P* < 0.001, Wilcoxon matched-pairs signed rank test was used in D. (**E**) Scatter diagram and quantification of the peak value difference between the R Ca^2+^ and Ω Ca^2+^, which was abolished in *nlp-18*, *ckr-1* and *nlp-18; ckr-1* mutants. Neuronal specific expression of NLP-18 in ASI in *nlp 18*, and CKR-1 in SMD in *ckr-1* could restore the Ω Ca^2+^ (n ≥ 13 animals). \*\*\**P* < 0.001, Mann-Whitney test. Data are expressed as mean ± SEM.

In *nlp-18* or *ckr-1* mutants, SMDD continued to exhibit oscillatory Ca^2+^ signals during reversal and turns similar to wild type animals (**Supplementary Figure S5A, B**). We found that the removal of *nlp-18* or *ckr-1* does not alter SMDD’s R Ca^2+^ peak and dynamics (**Supplementary Figure S5C-E**). We also did not observe elevation of R Ca^2+^ elevation even when animals performed shallow turning (**Figure 6D**; **Supplementary Figure S5A, B**; **Supplementary movie S4**). Consistent with the non-additive steering defects exhibited by *nlp-18; ckr-1* double mutant animals, SMD neurons exhibited a similar degree of reduction of turning-related elevation to either single mutant animals. In *nlp-18* mutant animals, expression of NLP-18 in ASI neurons restored Ω Ca^2+^ (**Figure 6D, E**). In *ckr-1* mutant animals, expression of CKR-1 in SMD also restored Ω Ca^2+^ (**Figure 6D, E**). Together, NLP-18-CKR-1 signaling promotes the Ω turn.

### CKR-1 is a receptor of NLP-18

Our functional data propose that NLP-18 and CKR-1 activate Gα_q_ signaling to promote robust escape steering. We investigate whether this effect is direct: CKR-1 as a receptor of NLP-18.

First, we tested the hypothesis in the heterologous *X. Laevis* oocyte system. Synthetic NLP-18 did not elicit endogenous currents in control oocytes (**Figure 7A**, empty vector). When *ckr-1* cRNA was injected in the oocyte, synthetic NLP-18 (NLP-18a to NLP-18e) evoked robust currents (**Figure 7A, B**). There are five predicted NLP-18 peptides, four with the hallmark C-terminal residues for the NLP peptides ‘FAFA’ (NLP-18b, c, e) or ‘FA’ (NLP-18a) (**Figure 7A**), and one without (NLP-18d). While all evoked currents, those with the FA motif were more potent (**Figure 7B**). Indeed, truncation of the FAFA residues from NLP-18c (NLP-18cΔ reduced current (data not shown), predicting a necessary motif for high-affinity binding. Application of five NLP-18 neuropeptides together (1 μ larger current than single peptides, indicating activation saturation (**Figure 7A, B**). Under this condition, the half-effect concentration (EC_50_) of NLP-18a was ∼13 nM (**Figure 7C**), within the range for cognate receptor activation (Rogers et al., 2003).

**Figure 7.**
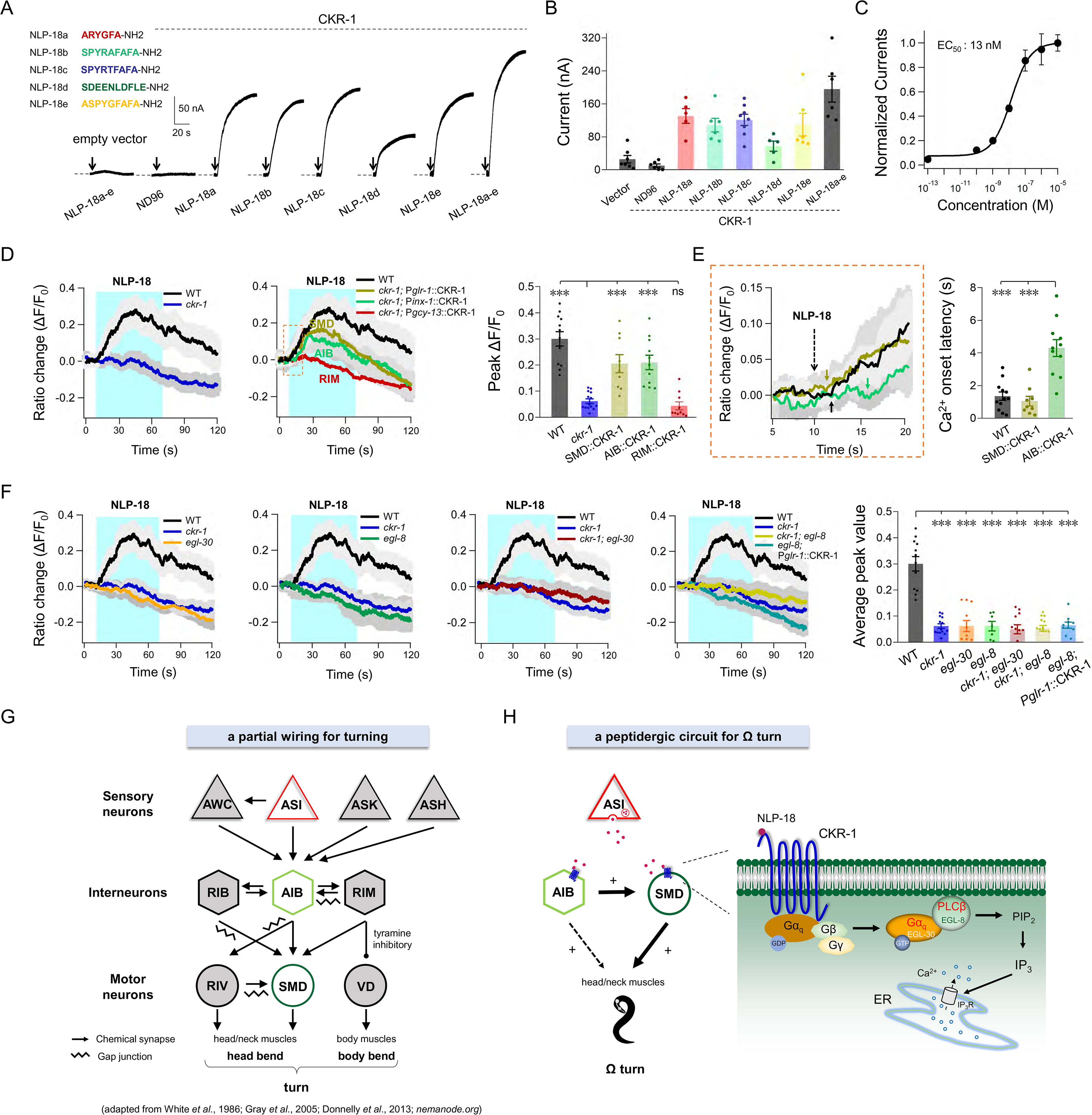
CKR-1 is a receptor of NLP-18. (**A**) Representative current traces evoked by different NLP-18 neuropeptides (1 μM) in the *Xenopus laevis* oocytes injected with CKR-1 cRNA. Five synthetic NLP-18 neuropeptides’ sequences were denoted by colors (red NLP-18a, green NLP-18b, blue NLP-18c, dark green NLP-18d, golden NLP-18e). Arrows indicate the puffing onset time. (**B**) Quantification of the NLP-18 evoked peak currents (n ≥ 5 oocytes). (**C**) The dose response of the NLP-18a evoked currents, revealing an EC_50_ of 13 nM fitted by hill equation. (**D**) NLP-18 neuropeptides (mixed NLP-18a-e, 1 μM each) evoked robust Ca^2+^ transient from the SMD neurons, which was terminated in the *ckr-1* mutant. The terminated Ca^2+^ transient could be rescued by expression of CKR-1 in SMD, AIB but not in RIM neurons (n ≥ 9 animals). ns, not significant, \*\*\**P* < 0.001, Mann-Whitney test. (**E**) A delayed activation occurred by expression of CKR-1 in AIB (n ≥ 9 animals). \*\*\**P* < 0.001, Mann-Whitney test. (**F**) SMD Ca^2+^ transients evoked by NLP-18 mixed neuropeptides (NLP-18a-e, 1 μM each), exhibits remarkable decrease in *egl-30*, *egl-8* single mutant and *ckr-1; egl-30*, *ckr-1; egl-8* double mutants. Expression of CKR-1 in SMD neurons could not rescue the Ca^2+^ response of *ckr-1* in *egl-8* mutant background (n ≥ 7 animals). \*\*\**P* < 0.001, Mann-Whitney test. (**G, H**) Working model. (G) Partial hard-wired neural circuit diagram for turning in *C. elegans*. (H) A neuropeptidergic circuit and signaling pathway for omega turn (escape steering): head bending motor neuron SMD receive NLP-18-CKR-1 neuromodulation inputs from ASI and may simultaneously receive neurotransmission inputs from AIB (+). AIB strengthen the head muscle activity through SMD as well as other neurons (dash line). Sensory neurons are represented by triangles, interneurons are represented by hexagons, motor neurons by circles. Arrows represent connections via chemical synapses, which may be excitatory or inhibitory. Wave lines represent connections by electrical synapses. Data are expressed as mean ± SEM.

Next, we asked whether NLP-18 could activate endogenous CKR-1 in *C. elegans* neurons. We applied a *C. elegans* dissection preparation to expose SMD motor neurons that expressed GCaMP6s (**Figure 6**), and assessed their calcium dynamics evoked by synthesized NLP-18 (NLP-18a-e mix) (see Methods). Application of synthetic NLP-18 evoked ∼30% increase at peak Ca^2+^ signals (**Figure 7D**). Evoked Ca^2+^ increase was abolished in *ckr-1* mutant animals, and were partially restored by CKR-1 expression in SMD (**Figure 7D**). Consistent with that CKR-1 encodes a Gα_q_ protein-coupled receptor, NLP-18 evoked Ca^2+^ increase in SMD was also abolished in *egl-30* and *egl-8* mutant animals (**Figure 7F**). In *ckr-1; egl-30* and *ckr-1; egl-8* double mutants, the Ca^2+^ transient increase was reduced to the same level in *egl-30* and *egl-8* single mutants, respectively. In the *ckr-1; egl-8* mutant, restoring CKR-1 expression in SMD could not rescue the Ca^2+^ increase (**Figure 7F**). We conclude that CKR-1 functions as a direct receptor of NLP-18 to increase SMD motor neuron’s activity by Gα_q_ signaling.

### AIB functions mainly through SMD

During escape, the rise and fall of AIB activity transits the animal from reversal to turn (Piggott et al., 2011; Wang et al., 2020), which led by a deep head bend that are contributed by SMD calcium increase. Expression of CKR-1 in AIB robustly rescued *ckr-1* mutant animal’s escape defects (**Figure 4C**). We found that restoring CKR-1 in AIB also rescued SMD’s Ca^2+^ increase (**Figure 7D**) but with an extended delay (latency 4.3±0.5 s) when compared to restoring CKR-1 in SMD (latency 1.1±0.3 s, *P* < 0.001) or wild type animals (latency 1.4±0.3 s, *P* < 0.001) (**Figure 7E**). This is consistent with AIB functioning at the upper circuit layer to activate turning (Gray et al., 2005).

The observation that other CKR-1-expressing interneuron such as RIM (P*gcy-13*::CKR-1) was not able to restore escape defects or SMD’s calcium increase suggests that NLP-18-CKR-1 signaling pathway may function through hard-wired circuit connections (**Figure 7D**). Indeed, upon ablation of SMD neurons by miniSOG, expression of CKR-1 in AIB severely reverted its rescuing effect in escape steering (**Supplementary Figure S6**). Interestingly, they maintained some capacity of full escape steering (**Supplementary Figure S6**), suggesting that AIB may function through a minor, secondary pathway to promote turning.

In summary, we show here that NLP-18, released mainly from the ASI sensory neurons, activate the CKR-1 GPCR in the SMD head motor neurons and AIB interneurons to strengthen escape steering (**Figure 7G, H**). The NLP-18-CKR-1 signaling pathway is an integral component of the escape response.

## Discussion

With increasingly detailed understanding of the molecular and cellular composition of *C. elegans* wiring (Bargmann, 1998; Cook et al., 2019; Taylor et al., 2021; White et al., 1986; Witvliet et al., 2021), it is possible to functionally dissect the role of neuropeptidergic signaling at single neuron resolution. Through a dedicated neural network, *C. elegans* generates an innate motor response to escape noxious stimuli. We discover a peptidergic (NLP-18)-GPCR (CKR-1) signaling pathway as a necessary component of the innate escape steering. Instead of a slow and lasting modulation of a circuit state, this signaling event functions selectively and temporally to enable reorientation steering, the omega turn.

### A necessity of peptidergic signaling in innate behaviors

Shared characteristics of innate behaviors reflects shared properties of the underlying circuits. Across species, escape behaviors are instinctual, executed with speed and robustness. The fast forms of neural transmission, through electrical and excitatory synaptic transmissions in the reflexive escape motor response, have been shown in vertebrate like larval zebrafish (Dunn et al., 2016; Miller et al., 2017).Withdraw, however, is only one motor step of the escape response. Previous and ongoing work has allowed *C. elegans* system to examine this response for not only other steps, but also their transitions. This enables a systematic investigation and modeling a robust, but not rigid behaviors from molecular, cellular and systems level. To date, the fast neural transmission that drive escape circuits for escape has been largely characterized (Chalfie et al., 1985; Gray et al., 2005; Pirri and Alkema, 2012; Wang et al., 2020).

Working in parallel with neurotransmitters, neuropeptides exert effects for innate behavior on relatively slow timescales through intracellular signaling cascades (Bargmann, 2012; Bargmann and Marder, 2013). For example, the neuropeptides hypocretin/orexin produced in hypothalamic neurons, are crucial regulators of sleep and wakefulness (Sakurai, 2007). Here, we identified a peptidergic signaling pathway that underlies a motor behavior during escape. The NLP-18-CKR-1 signaling pathway specifically enhances escape steering, but not in other modalities such as spontaneous velocity. This functional specificity of NLP-18 reflects spatial specificity by origin (ASI) and target neurons (AIB and SMD), as well as the temporal specificity by its activation of head motor neurons during turning. Such a mode of function differs from what generally sought to be a slow, long range, and long-lasting form of neuromodulation.

Consistent with this notion, NLP-18-CKR’s requirement for escape steering was independent of the type of stimuli or the environmental food signals. Because SMD motor neurons exhibit normal calcium dynamics during spontaneous backward movement in *nlp-18* and *ckr-1* mutants, these results support the notion that NLP-18 secretion occurs specifically during escape to regulate SMD and AIB for Ω turn. However, how this release is regulated remains unknown and requires further investigation.

### Functional conservation of GPCR receptors by sequence-divergent signaling peptides

CKR-1 exhibits high sequence homology to the vertebrate CCK1R (or CCKAR) family GPCRs, and same as the human CCK1R (Dufresne et al., 2006), is a Gα_q_ protein-coupled receptor. Human CCK1R is expressed in the gastrointestinal tract and discrete brain areas, coinciding with CKR-1’s expression in *C. elegans* intestine and a selected group of sensory, motor- and inter-neurons. These similarities implicate potential common physiological contributions. Activation of CCK1R induces satiety, an innate feedback on food intake (Jensen, 2002) and disruption of human CCK1R causes psychiatric disorders (Shintaku et al., 1992). *C. elegans* NLP-18-CKR-1 signaling involves the gustatory sensory neuron and escape response. NLP-18’s primary neuronal source ASI are known to secrete hormone-like peptides insulin and TGF-beta. With only dense core vesicles, ASI exhibits the classic morphology of secretory neurons (Witvliet et al., 2021). There might be potential conservation of CKR-1’s role in neural circuits.

Neuropeptides are generally considered to be species-specific due to low primary sequence identity. Our investigation of CKR-1’s physiological role led to its cognate ligand NLP-18, challenging this notion. There is no steering defect in loss-of-function of *ckr-2*, another putative CCK homologue, supporting NLP-18’s specificity for CKR-1. Known CCK1R ligands, CCK in vertebrate, gastrin in arthropod, and NLP-18 in *C. elegans* do not exhibit strong sequence similarity. However, CCK was also found in the gastrointestinal tract (Ivy and Oldberg, 1928) and the brain (Beinfeld, 1983), bearing resemblance NLP-18’s presence in the intestine and sensory and interneurons. Both peptides show dose-dependent activation of the CKR receptor coupled with G□ signaling. NLP-18 exhibits high affinity to CKR-1, with initiation activation at 1 nM and half-activation at 13 nM. CCK-8 was reported to activate CCKR1 between 1 nM to 10 nM, and peak activation at 100 nM (Shigeri et al., 1996). NLP-12 was considered to be CCK-8’s *C. elegans* homolog by sequence (Janssen et al., 2008), but *nlp-12* mutant animals don’t exhibit escape steering defect (**Supplementary Figure 1B**). These findings argue for assessing physiological homology of peptidergic signaling by receptors, instead of the primary sequence of ligands. Identifying NLP-18 as a cognate ligand further expands the search of multiple ligands for CCK receptors.

### Peptidergic signaling functions through a hard-wired circuit

NLP-18’s requirement for escape steering is not specific to the type stimuli. From mechanical (head touch), chemical (copper), to high osmolarity (glycerol), all required NLP-18 to evoke deep Ω turns. Similarly, its source, the ASI sensory neurons, are not the primary sensory receptors any of these stimuli. NLP-18 and ASI form an integral component of the escape circuit.

Primary nociceptor neurons make direct and indirect input onto the layers of interneurons, premotor interneurons, and motor neurons of the *C. elegans* escape circuit. The steering behavior is regulated by sensory neurons, such as ASI (**Figure 7G**) that may not directly evoke escape responses. ASI are gustatory and multimodal, known for their roles in food-related modulation, including insulin-related chemotaxis behaviors and dauer formation (Bargmann and Horvitz, 1991). In the absence of food, ASI suppress Ω turn frequency by inhibiting AWC, an olfactory nociceptor (Gray et al., 2005) and other nociception (Guo et al., 2015). This differs from the finding here that ASI is required for and enables steering by NLP-18-dependent increase of head motor neuron activity. We speculate that ASI play dual roles: they are required for escape steering that is independent from food-sensing, but can activate a pathway to suppress Ω turn to allow food searching when animals are hungry.

NLP-18’s target neurons are similarly specific. The head motor neurons SMD’s activity level determines Ω turn frequency (Gray et al., 2005),head-bend amplitude during foraging (Shen et al., 2016; Yeon et al., 2018) and post-reversal turn amplitude (Kaplan et al., 2020). Specific loss of elevated Ω *1* mutants demonstrates the requirement of peptidergic signaling. Intriguingly, depletion of CKR-1 in either SMD or AIB reduces escape steering. AIB interneurons transit animals from reversal to turn (Gray et al., 2005; Wang et al., 2020). By receiving chemical synapses from multiple sensory neurons including ASI, they send chemical and electrical synapses to motor neurons SMD. Because restoration of CKR-1 in AIB alone partially restored the escape steering behavior and NLP-18-evoked SMD Ca^2+^ amplitude with a delay, AIB may promote escape steering through chemical synaptic activation of SMD (**Figure 7H**). ASI have no direct wiring to SMD, NLP-18-CKR-1 pathway may employ paracrine or endocrine signaling combined with hard-wired neuronal circuit for escape modulation. We propose that NLP-18-CKR-1 signaling strengthens the functional connection of the ASI-AIB-SMD circuit for robust escape steering.

RIM interneurons form a subcircuit with AIB and they exhibit coordinated activity changes during reversal (Kawano et al., 2011). Ablation of RIM suppresses turnings during thermal taxis (Ji et al., 2021), but unlike the case of ASI ablation, *nlp-18* and *ckr-1*, RIM’s effect may be more context dependent. Although both *nlp-18* and *ckr-1* are expressed in RIM, depletion or overexpression of NLP-18 or CKR-1 in RIM did not change or rescue turning phenotypes of wildtype, *nlp-18* or *ckr-1* mutants. Co-expression of NLP-18 in both ASI and RIM did achieve a slight improvement in behavior rescue compared to ASI along, although we cannot rule out an overexpression effect. The lack of rescuing effect on its own may reflect its limited capacity for secreting NLP-18: unlike ASI, RIM exhibit mostly clear synaptic vesicles with a small number of dense core vesicles (Witvliet et al., 2021), supporting an implication from the expression study that RIM is primary a glutamatergic neuron (Alkema et al., 2005).

In summary, we identified a temporally and spatially controlled neuropeptidergic neuronal signaling that function with and might through a hard-wired circuit to drive escape steering. The head motor neuron that codes for turn angles requires this signaling to execute robust steering. With a specific input from a gustatory neuron to potentiate the interneurons and motor neurons that encode turning (**Figure 7H**), this wiring addition potentially renders food state-dependent modulation of navigation.

## Materials and Methods

### Constructs, transgenic arrays and strains

The complete lists of constructs, transgenic lines and strains generated or acquired for this study are provided in Supplementary Table 1-3. All *C. elegans* strains were cultured on the standard Nematode Growth Medium (NGM) plates seeded with OP50 and maintained at 22 °C. Unless otherwise stated, the wild type animal refers to the Bristol N2 strain.

### Molecular biology

All expression plasmids in this study were constructed by Three-Fragment Multisite gateway (Invitrogen, Thermo Fisher Scientific, Waltham, MA, USA) (Magnani et al., 2006). Three-Fragment Multisite gateway system consists of three entry clones. Three entry clones comprising three PCR products (promoter, gene of interest, *sl2d-*GFP, *sl2d*-wCherry or *unc-54*-3’UTR, in name of slot1, slot2 and slot3, respectively) were recombined into the *pDEST-R4-R3*, *pDEST-R4-R3-unc-54* 3’UTR, *pBCN44-R4R3-Plin-44*::GFP and *pBCN44-R4R3-Pvha-6*::GFP destination vectors by using standard LR recombination reactions to generate the expression clones.

All promoters used in this study were generated by PCR against a mixed-stage N2 *C. elegans* genomic DNA. Promoters’ genomic sequences were used to substitute for the *rab-3* fragment in standard BP reaction-generated entry clone A with the In-Fusion method, using ClonExpress® One Step Cloning Kit.

To generate the entry clones slot2 and slot3, we used standard BP recombination reactions. An entry clone B contributing sequences of slot 2 in the expression plasmid contains sequence of a target gene, i.e., *nlp-18*, *ckr-1*, or any gene of interest. The “B” entry clones were constructed by using of *pDONR™ 221* donor vector (Invitrogen) through BP reactions. All the DNA fragments used to construct entry clone B involved in this project were amplified by PCR with the primers containing attB1 and attB2 recombination sites.

An entry clone C contributing sequences of slot 3 in the expression plasmids contains a sequence of *unc-54*-3’UTR, *sl2d-*GFP or *sl2d-*wCherry. The “C” entry clones were constructed by use of *pDONR-P2R-P3* donor vector through standard BP reactions. The corresponding PCR products with attB2R and attB3 sites were amplified by PCR. The “C” entry clones containing *unc-54*-3’UTR, *sl2d-*GFP and *sl2d-*wCherry were used to construct expression plasmids.

### ZIF-1-ZF1 system

ZIF-1-ZF1 system was used to degrade target protein in specific tissue and neurons respectively (Armenti et al., 2014). In ZIF-1-ZF1 system, P*ckr-1*-ZF1-SL2-NLS-GFP was knocked in before termination codon of *ckr-1* using CRISPR-Cas9 to generate GB01 *gaaIs1* [P*ckr-1*-ZF1-SL2-NLS-GFP] strain. Using different promoter combined ZIF-1 and *sl2d-*GFP or *sl2d-*wCherry by LR reaction to construct expression plasmids then co-injecting P*lin-44*::GFP into GB01 to degrade CKR-1 protein in specific neuron.

Transgenes arrays and strains Transgenic animals that carry non-integrated, extra-chromosomal arrays (*gaaEx*) were generated by co-injecting an injection marker with one to multiple DNA construct at 5-30 ng/μl. Animals that carry integrated transgenic arrays (*gaaIs*) were generated from the *gaaEx* animals by UV irradiation, followed by outcrossing against N2 at least 4 times. L4 stage or young adults (24 hr post L4 stage) hermaphrodites were used in all experiments. Other genetic mutants used for constructing transgenic lines and compound mutants were obtained from the *Caenorhabditis Genetics Center* (*CGC*).

### Neuronal manipulation

To dissect the role of ASI neurons in locomotion modulation, we expressed mito-miniSOG into these neurons driven by P*gpa-4*. Ablation of ASIs was performed using a homemade LED box, where the standard NGM were exposed under a homemade 470 nm blue LED light (8.3 mW/cm^2^) for 30-45 minutes. To monitor the specificity and efficacy of cell ablation, cytoplasmic wCherry was co-expressed with miniSOG (tomm-20-miniSOG-SL2-wCherry) in targeted neurons by the same promoter. Ablation was performed when animals were in the L2-L3 stage, which would be examined whether the wCherry fluorescence was absent after 24 hr. Late L4 or young adult stage animals were recorded for behavioral analyses.

For chemogenetic silencing the ASI neurons, we used neuronal specific promoter P*gpa-4* to drive expression of the Drosophila *HisCl* gene in the ASI neurons. To make NGM-HA plates, histamine (Sigma Aldrich histamine-dihydrochloride, 1 M stock in water) was added to NGM agar at ∼65°C. Histamine-free control plates were poured from the same NGM batch. NGM plates with 10 mM histamine were used to test the behavioral effects.

### Behavioral Analyses

Harsh head touch was delivered with a platinum wire pick. One hour prior to testing, normal cultured young adult hermaphroditic animals (8-12 hr post L4 stage) were transferred to plates with thin layer of OP50 bacteria. The worm head was touched with the edge in a top-down manner with a force of 100-200 μN (Li et al., 2011). For each strain, at least 30 worms were recorded in each group and experiment was repeated for 3-5 times. To avoid the possible stimulation-induced adaptation, each worm was tested for one time. The full omega was defined as the head contact with the body when animal turning after reversal, or else it was classified into shallow omega. A part of animals (2.7±0.4%) did not execute the omega turn but only reversal after stimulation. The proportion of full omega, shallow omega and only reversal was quantified to present the robustness of escape behavior. Omega curvature analysis utilized image J (National Institutes of Health) and Matlab (MathWorks). Extracting image which worm started to enter turning and showed the largest bending of head. Images from each animal were divided into 50 body segments for curvature analysis. The curvatures at each point along the worm centerline *k(s)* can be calculated with the coordinate of each point (*x(s), y(s)*) using the formula 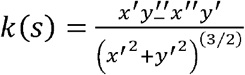 where s is the normalized location along centerline (head = 0, tail = 1), and the unit of *k* is pixel\(−1). Then *k* is normalized with the length of worm body *L*, resulting in the dimensionless *k∼(s)* using the formula 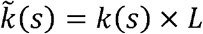. The curvature from 0%-60% body segment was plotted and converted as a color map. Omega angle was defined as the intersection angle from middle point of body (the 26^th^ cross line) to the closest points anterior and posterior of the animal. Thus, the angle of full omega is zero degree.

A single young adult hermaphrodite (12-18 hr post L4 stage), maintained on standard culture conditions, was transferred to a 60 mm imaging plate seeded with a thin layer of OP50. One minute after the transfer, a three-minute video of the crawling animal was recorded on a modified stereo microscope (Axio Zoom V16, Zeiss) with a digital camera (acA2500-60um, Basler). Post-imaging analyses utilized an in-house written MATLAB script. The central line was used to track. Images for velocity analysis from each animal were divided into 33 body segments. The mid-point was used to calculate the velocity and direction of movements between each frame.

Imaging plates were prepared as follows: a standard NGM plate was seeded with a thin layer of OP50 12-14 hr before the experiment. Immediately before the transfer of worms, the OP50 lawn was spread evenly across the plate with a sterile bent glass rod. All images were captured with a 10× objective at 10 Hz.

### Confocal Fluorescence Microscopy

L4 stage transgenic animals expressing fluorescence markers were picked a day before imaging. Worms were immobilized by 2.5 mM levamisole (Sigma-Aldrich) in M9 buffer. Fluorescence signals were captured from live worms using a Plan-Apochromatic 60× objective on a confocal microscope (FV3000, Olympus) in the same conditions.

### Calcium imaging

The free-tracking Ca^2+^ imaging system include two independent modules: behavior tracking system and fluorescence recording system. Worm behavior was imaged under dark-field illumination in the near-infrared (NIR). Pick a worm on 6 cm NGM plates and then put it on a THOPLABS spw602 XY motorized stage controlled. Imaging plates were prepared according to the behavior imaging plates. Tracking imaging was conducted using a 10× inverted objective (Olympus, Japan) and recorded using a CCD camera (Point Grey Research, CM3-U3-13S2M, Canada). Custom real-time computer vision software kept the worm centered in the field of view via tracing centerline of worm. Fluorescence recording system records neuronal Ca^2+^ activity by using sCMOS digital camera (Hamamatsu, Japan) with 20× objective (Olympus, Japan). To simultaneously image wCherry and GCaMP6 side-by-side, we used a two-channel imager (W-VIEW GEMINI, Japan). Red and green channel fluorescent images were recorded simultaneously at 8 fps with 10 ms and 70 ms exposure time, respectively. Fluorescent images were captured using HCImage software (Hamamatsu). Neural activity was reported as normalized deviations from baseline of the ratio between GCaMP6s and wCherry fluorescence, 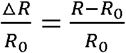, where 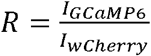. The baseline *R_0_* is defined as the minimum of *R*. The intensities *I_GCaMP6s_* and *I_wCherry_* were measured as the pixel intensity in the green and red channels, respectively. Custom MATLAB scripts were used to extract the pixel intensity for each frame. We focused on SMDD for quantitative analyses was that in our reporter, SMDD was isolated from other neurons when compared to SMDV, making it easier to extract high-quality activity traces. On a side note, we noticed that when we performed calcium imaging recordings, it was not difficult to find animals executing dorsal Ω turns, which occurs much less frequently under regular light microscopy.

For NLP-18 puffing evoked Ca^2+^ imaging, worms were glued and dissected to expose SMD neuron as described for electrophysiological recording. Then imaged with a 60× water objective (Nikon, Japan) and sCMOS digital camera (Hamamatsu ORCA-Flash 4.0 V2, Japan) at 10 Hz with expose time 100 ms. For all the SMD puffing neuropeptides experiments, we recorded the neuronal Ca^2+^ transient by the perfusion of the NLP-18 neuropeptides for 60 seconds and washing out for 50 seconds by normal bath solution. Ca^2+^ transients’ onset latency was calculated by the time difference from the perfusion onset to Ca^2+^ initiation. The Ca^2+^ transients of SMD soma were analyzed by Image-Pro Plus 6 (Media Cybernetics, Inc., Rockville, MD, USA).

### Oocytes expression and electrophysiology

CKR-1 expression in *X. laevis oocytes*: CKR-1 cDNAs were flanked between BamHI and HindIII sites by PCR and cloned into the *pGH19* vector. CKR-1 cRNAs were prepared using the mMessage Machine kit (Ambion). *X. laevis* oocytes were injected with 50 ng of CKR-1 receptor sense cRNAs. Injected oocytes were then incubated at 18°C in the ND96 medium for 2-3 days before recording.

Current recordings were made using the two-electrode voltage-clamp technique at a holding potential of –80 mV as described (Rogers et al., 2003). Oocytes were continuously superfused with ND96 solution contains: 96 mM NaCl, 2.5 mM KCl, 1 mM MgCl_2_ and 5 mM HEPES, pH 7.3. The recording chamber was perfused with high-K^+^ solution to reverse the K^+^ gradient (Rogers et al., 2003) and measured the ligands (NLP-18a-e) dependent outward Ca^2+^-gated chloride currents (Oron et al., 1985). The pipette solution contains 3M KCl. The recording high-K^+^ bath solution contains: 96 mM KCl, 2.5 mM NaCl, 1 mM MgCl_2_ and 5 mM HEPES, pH 7.3. Peptide perfusion was terminated by washout with high-K^+^ solution and subsequent switching to ND96 solution. Data were acquired with Clampex 8.0 software (Molecular Devices) and analyzed offline with Clampfit (Molecular Devices). Peptides were synthesized by the Guoping Pharmaceutical Company (Hefei, Anhui Province, China).

### Statistical analysis

The Mann-Whitney test, two-tailed Student’s *t*-test, Two-way analysis of variance (ANOVA) tested were used to compare data sets. Statistical analysis of proportion was performed with the Fisher’s exact test. Kolmogorov-Smirnov was used analysis shallow exit Ω angle cumulative fraction. The paired Ca^2+^ values from same SMDD neurons were analyzed by Wilcoxon matched-pairs signed rank test. *P* < 0.05 was considered to be statistically significant (\**P* < 0.05, \*\**P* < 0.01, \*\*\**P* < 0.001). Graphing and subsequent analysis were performed using Igor Pro (WaveMetrics), Clampfit (Molecular Devices), Image J (National Institutes of Health), Matlab (MathWorks), GraphPad Prism 8 (GraphPad Software Inc.), and Excel (Microsoft). Phylogenetic tree was analyzed by MEGA 6.60. For behavior analysis, electrophysiology and fluorescence imaging, unless specified otherwise, each recording trace was obtained from a different animal. Data were presented as the mean ± SEM.

## Author Contributions

L.C. and S.G. conceived experiments, analyzed data and wrote the manuscript. Y.L., P.S., Y.W. and Z.Y. performed experiments and analyzed data. W.H., H.L., M.G., Z.W., Y.Z., P.F., L.C., L.T. and H.M. contributed to the experiments. M.Z. provided unpublished reagents, assisted with experimental design and edited the text.

## Acknowledgements

We thank Lijun Kang, Quan Wen, and Jianfeng Liu for comments and discussion. This research was supported by the Major International (Regional) Joint Research Project (32020103007), the National Natural Science Foundation of China (31871069), and the Overseas High-level Talents Introduction Program, and NSFC grant (31771294) to L.C. We thank *Caenorhabditis Genetics Center*, which is funded by the NIH Office of Research Infrastructure Programs (P40 OD010440), for strains.

---

*Figure S1. Genetic screening and different stimulations.*

(**A**) The full Ω proportion was decreased in carboxypeptidase E *egl-21* mutant and dense-core vesicle priming required CAPS (calcium-dependent activator protein for secretion) ortholog *unc-31* mutant, but was not changed in insulin pathway genes *daf-2* and *daf-16* (n ≥ 4 trials). ns, not significant, \*\*\**P* < 0.001, Student’s *t*-test. (**B**) The full Ω proportion was significantly reduced in *nlp-18* mutant from the screening in a group of neuropeptide-like peptide genes (n ≥ 3 trials). ns, not significant, \*\*\**P* < 0.001, Student’s *t*-test. (**C**) Quantification of the full Ω proportion in a group of FMRFamide-related peptides genes, from which *flp-1, 18, 20* exhibit significant defect (n ≥ 3 trials). ns, not significant, \**P* < 0.05, \*\*\**P* < 0.001, Student’s *t*-test. (**D**) Propensity, *entry* Ω curvature and shallow *exit* Ω angle evoked by chemical stimulation (100 mM Cu^2+^) in wild type and *nlp-18* mutants (n ≥ 3 trials). (**E**) Propensity, *entry* Ω curvature and shallow *exit* Ω angle evoked by osmotic stimulation (1M Glycerinum) in wild type and *nlp-18* mutants (n ≥ 3 trials). Each trial with at least 30 worms were tested. Statistical analysis of proportion was performed with the Fisher’s exact test. The *enter* Ω curvature was analyzed by Two-way ANOVA test. Kolmogorov-Smirnov was used to analyze shallow *exit* Ω angle cumulative fraction. \**P* < 0.05, \*\*\**P* < 0.001. All data are expressed as mean ± SEM.

*Figure S2. Sequence alignment and additional data of expression and rescue of nlp-18.*

(**A**) NLP-18 amino acid sequence alignment with other species. Identical amino acids are highlighted in color. (**B**) The co-localization of *nlp-18* in RIM and FLP neurons. The GFP driven by endogenous *nlp-18* promoter was co-localized with RFP driven by P*gcy-13* in RIM neuron and by P*mec-3* in FLP neuron, respectively. Arrows represent the worm orientation, A anterior, D dorsal. Scale bar, 20 μm. (**C, D**) The *entry* Ω curvature and shallow *exit* Ω angle defects in *nlp-18* mutant were moderately restored by the expression of NLP-18 in ASI + FLP and ASI + RIM neurons, and was fully restored when expressing NLP-18 in ASI + RIM + FLP neurons but was not in neurons without ASI. Two-way ANOVA test for *entry* Ω curvature in C, Kolmogorov-Smirnov test for shallow *exit* Ω angle cumulative fraction in D. Statistical significance is indicated as follows: ns, not significant, \**P* < 0.05, \*\**P* < 0.01, \*\*\**P* < 0.001. Data are expressed as mean ± SEM.

*Figure S3 ckr-1 exhibits same escape defects as nlp-18.*

(**A**) Instantaneous velocity in animals of respective genotypes. All mutants exhibited a comparable propensity and velocity as wild type N2 (n ≥ 13 animals). Data are expressed as mean ± SEM. ns, not significant, Two-way ANOVA test. (**B**) Propensity of directional movements in animals of respective genotypes, quantified by the animal’s mid-point displacement (n ≥ 11 animals). A movement of the centroid < 1 pixel/second is defined as pausing. Fisher’s exact test was used. (**C**) Screening of the candidate neuropeptide receptor mutants reveals that *ckr-1* dampens the full Ω proportion (n ≥ 4 trials). Data are expressed as mean ± SEM. \*\*\**P* <0.001, Student’s *t*-test. (**D**) Putative homologues of CKR-1 (with NCBI accession number) identified by blast search. Sequence alignment of 421 amino acids of CKR-1 using the Vector NTI Advance 11. Identical amino acids are highlighted in yellow, and similar ones are highlighted in grey. Seven predicted transmembrane regions are denoted by black boxes. The blue lines denote the deletion residues of *ok2502* allele used in this study, which truncated CKR-1 from 96 to 293 amino acid. (**E**) Propensity of head touch evoked escape responses (no Ω, shallow Ω and full Ω) in wild type, *nlp-18*, *ckr-1* and *nlp-18; ckr-1* double mutants in the absence of food (≥ 3 trials, at least 30 animals each trial). Fisher’s exact test was used. (**F**) Quantification of the *entry* Ω curvature in wild type, *ckr-1* and *nlp-18; ckr-1* mutants (n ≥ 100 animals). \*\*\**P* < 0.001, Two-way ANOVA test. (**G**) *ckr-1* and *nlp-18; ckr-1* show similar increased shallow *exit* Ω angle phenotype of *nlp-18* (n ≥ 45 animals) Kolmogorov-Smirnov test in G *(Left),* Mann-Whitney test in G *(Right),* \*\*\**P* < 0.001. Data are expressed as mean ± SEM.

*Figure S4. ckr-1 is functional in SMD and AIB neurons.*

(**A, B**) Additional evidence of neuronal specific expression of CKR-1 in different neurons in the *ckr-1* mutant. The *entry* Ω curvature and shallow *exit* Ω angle in the *ckr-1* mutant were restored by the expression of CKR-1 in AIB and SMD neurons, but not other neurons. All data are expressed as mean ± SEM. Two-way ANOVA test were used to access the statistical difference in A. Shallow exit Ω angle cumulative fraction was tested by Kolmogorov-Smirnov in B. Statistical significance is indicated as follows: ns, not significant, \*\**P* < 0.01 and \*\*\**P* < 0.001 in comparison with that of as indicated. Data are expressed as mean ± SEM.

*Figure S5. The Ω Ca^2+^ dynamics of SMD neurons was not affected*

(**A**) Representative images showing fluorescence of GCaMP6s and wCherry in the SMDD soma of P*glr-1*::GCaMP6s::wCherry transgenic worms during reversal and shallow Ω. (**B)** A representative Ca^2+^ transient trace of SMDD neurons color map of head bending in the process of reversal and turn from a free-behaving animal from a free-behaving animal during a shallow Ω turn. (**C**) The definition of the Ca^2+^ transient dynamics. (**D**) The raise time (10-90%) of the R Ca^2+^ transients and Ω Ca^2+^ transients showed no difference in all genotypes (n ≥ 13 animals). ns, not significant, Student’s *t*test. (**E**) Quantification of the peak value of R Ca^2+^ in *nlp-18*, *ckr-1* and *nlp-18; ckr-1* mutants and rescue strains. ns, not significant, Two-way ANOVA test. Data are expressed as mean ± SEM.

*Figure S6. AIB regulates escape steering partially through SMD*

(**A**) Composition of the head touch induced escape responses (no Ω, shallow Ω and full Ω) with or without SMD ablation (illuminated by blue light) in the AIB rescued or norescued transgenic strains (AIB::CKR-1) in *ckr-1* mutant (≥ 3 trials, at least 30 animals each trial). Statistical analysis of proportion was performed with the Fisher’s exact test. (**B, C**) Distributions of the *entry* Ω curvature and shallow *exit* Ω angle in the same conditions. Two-way ANOVA test in B. Kolmogorov-Smirnov was used in C. ns, not significant, \*\**P* < 0.01, \*\*\**P* < 0.001. Data are expressed as mean ± SEM.

***Supplementary Movie S1***: Head touch evoked a robust escape in wild type animals.

***Supplementary Movie S2***: Head touch evoked a robust escape in *nlp-18* mutants.

***Supplementary Movie S3***: R Ca^2+^ and Ω Ca^2+^ signals of SMDD neurons in free escaping wild type animals. SMDD neurons are circled.

***Supplementary Movie S4***: R Ca^2+^ and Ω Ca^2+^ signals of SMDD neurons in free escaping *nlp-18* mutant animals. SMDD neurons are circled.

