## Supplementary figures and images for "Escape Steering by Cholecystokinin Peptidergic Signaling"

### Figure 1

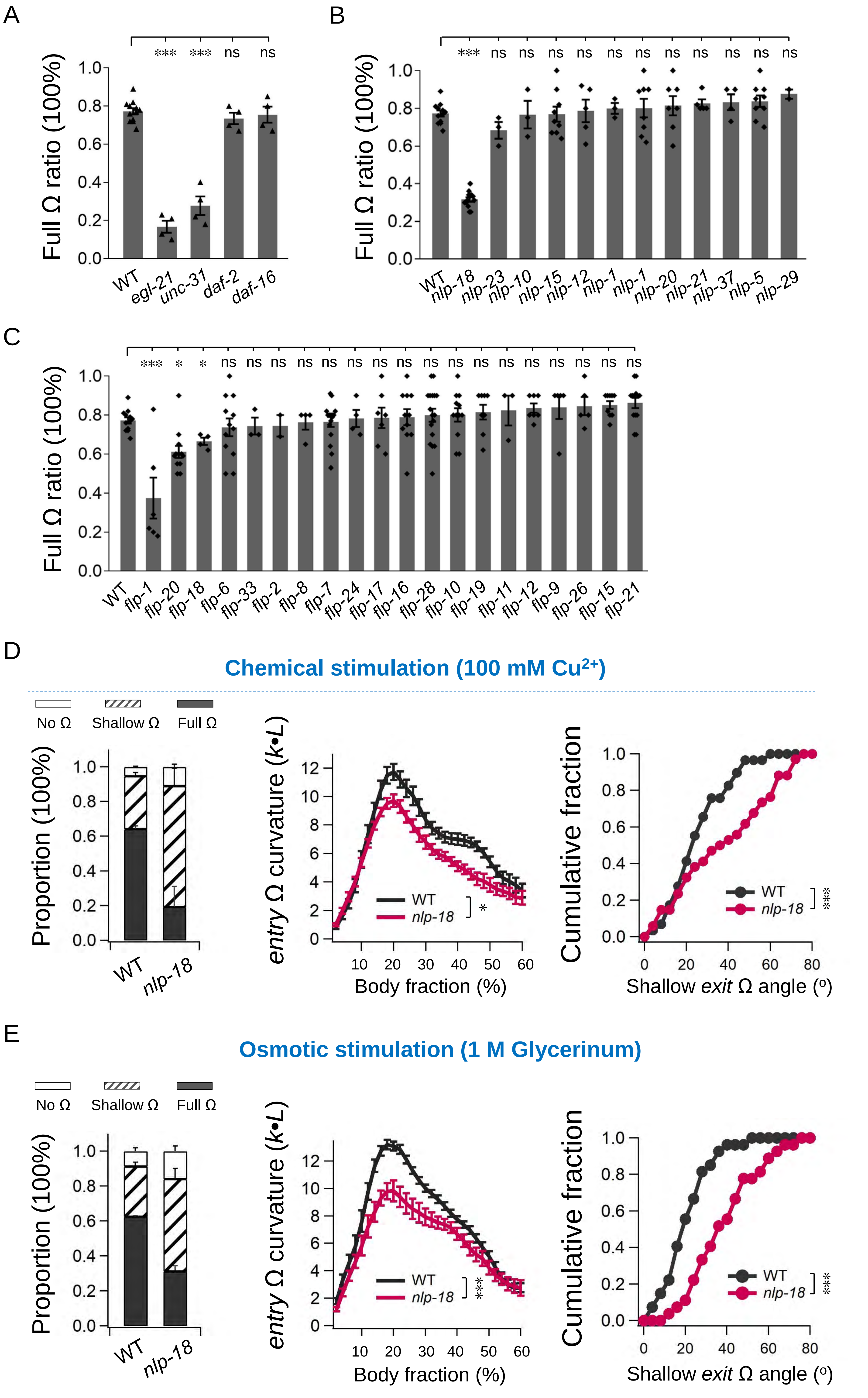

### Figure 2

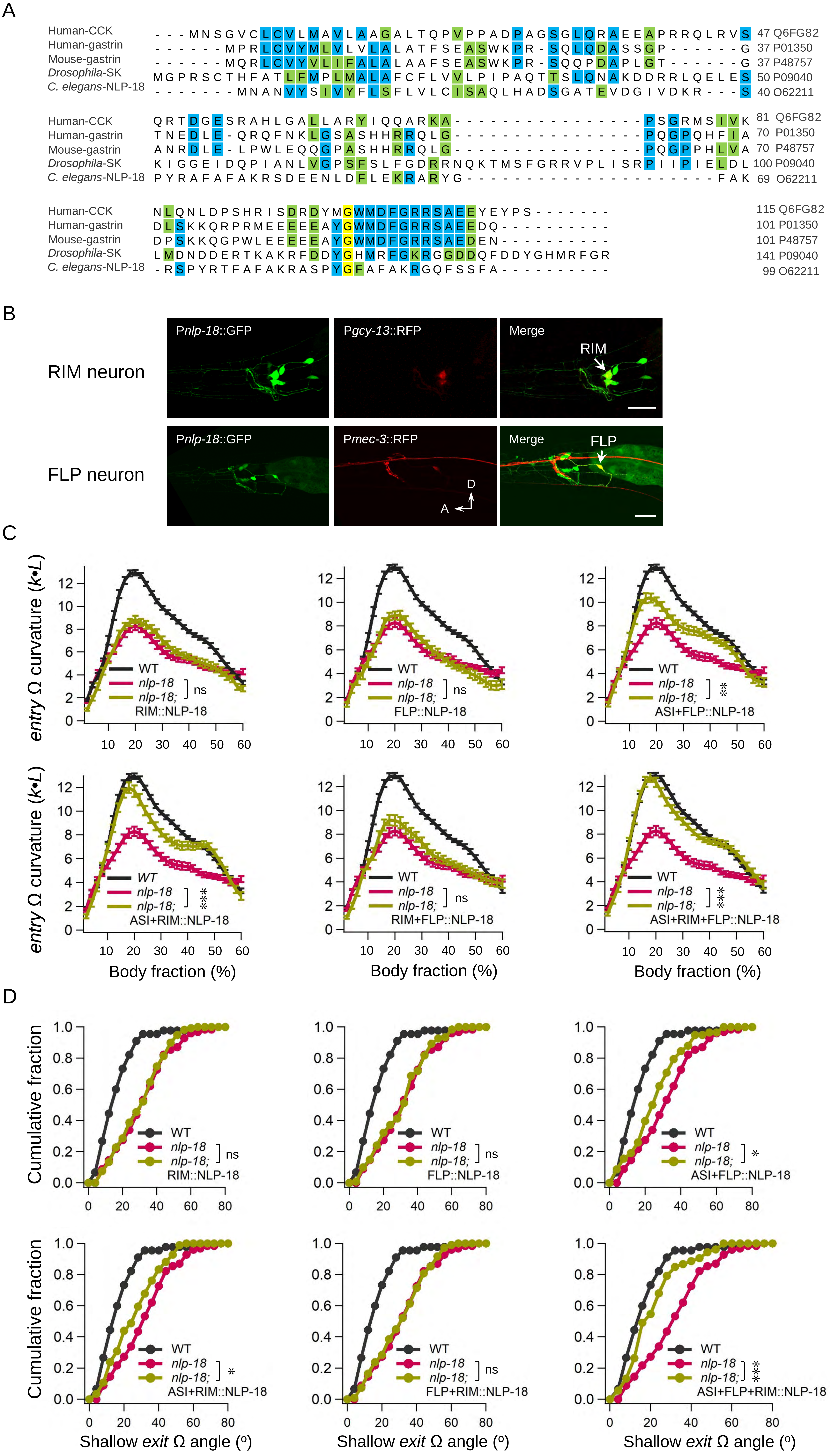

### Figure 3

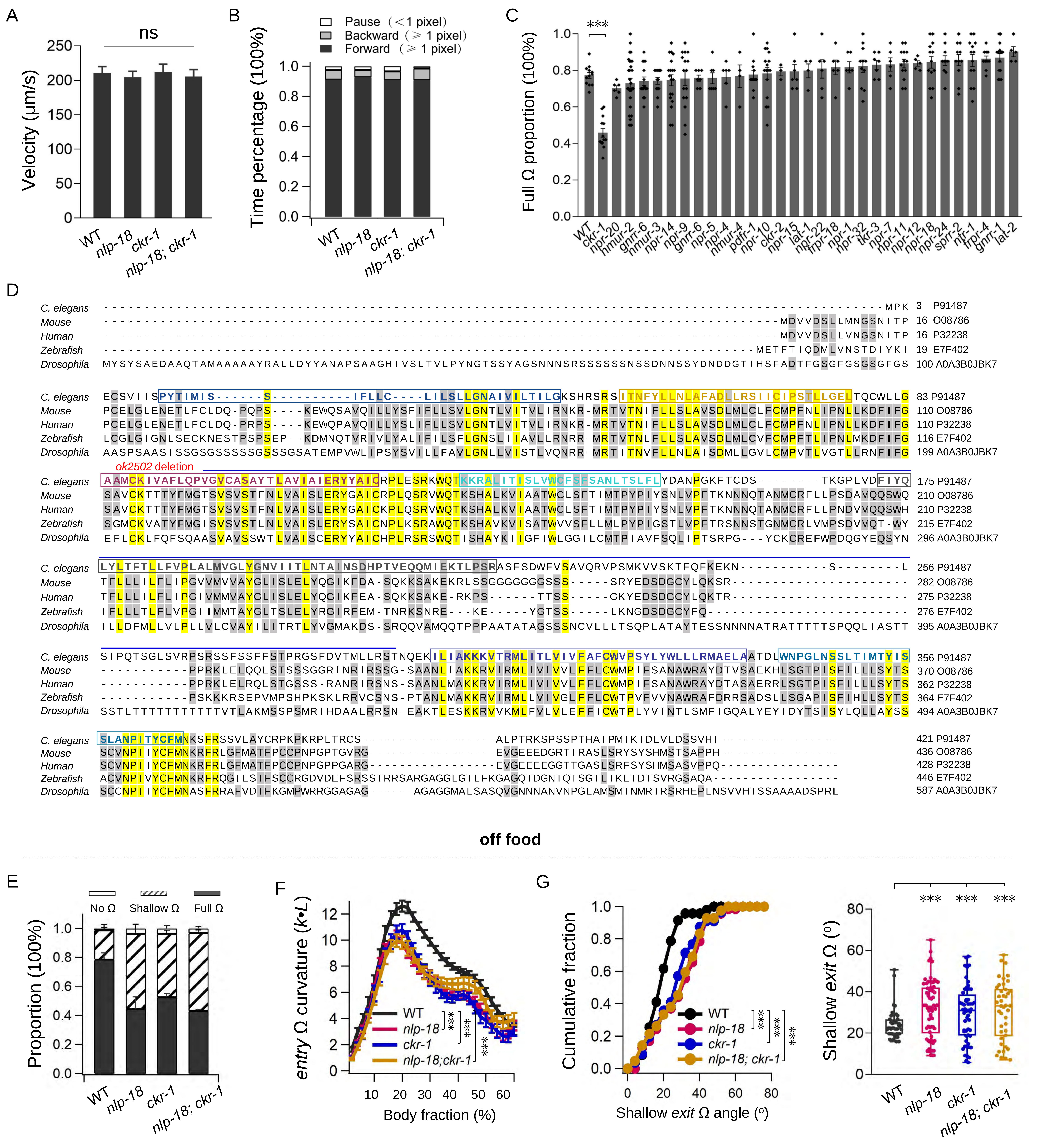

### Figure 4

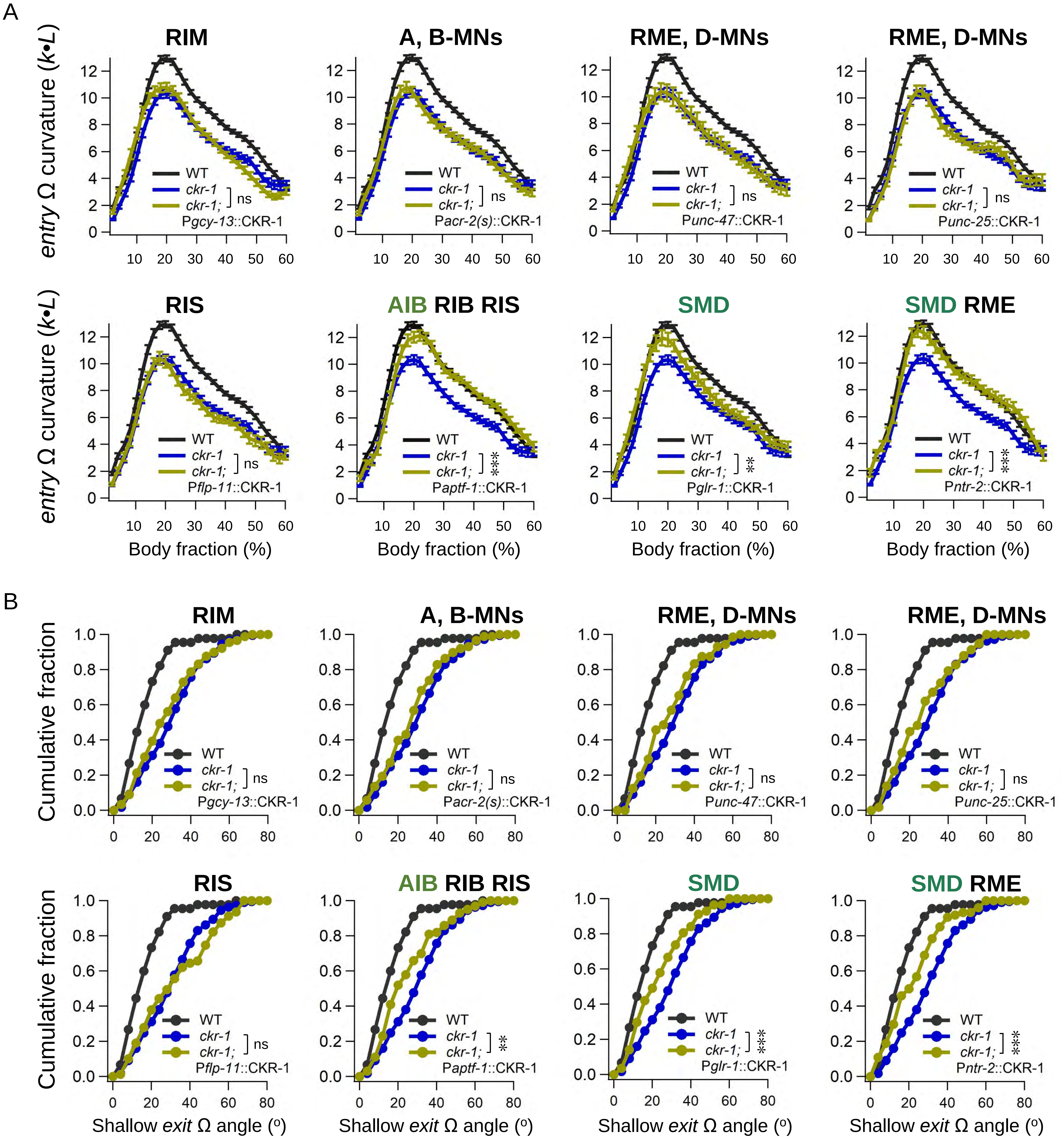

### Figure 6

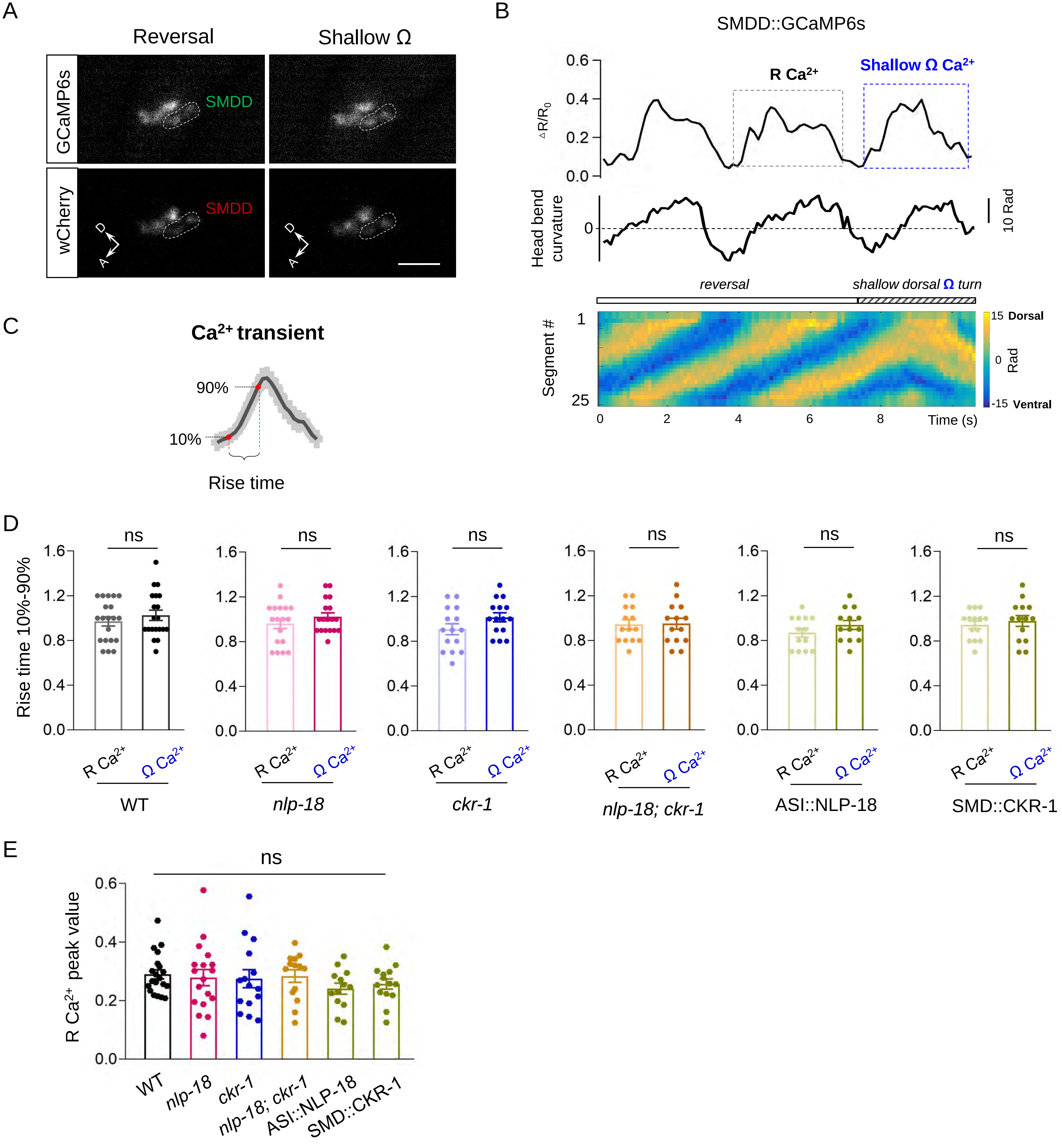

### Figure 7

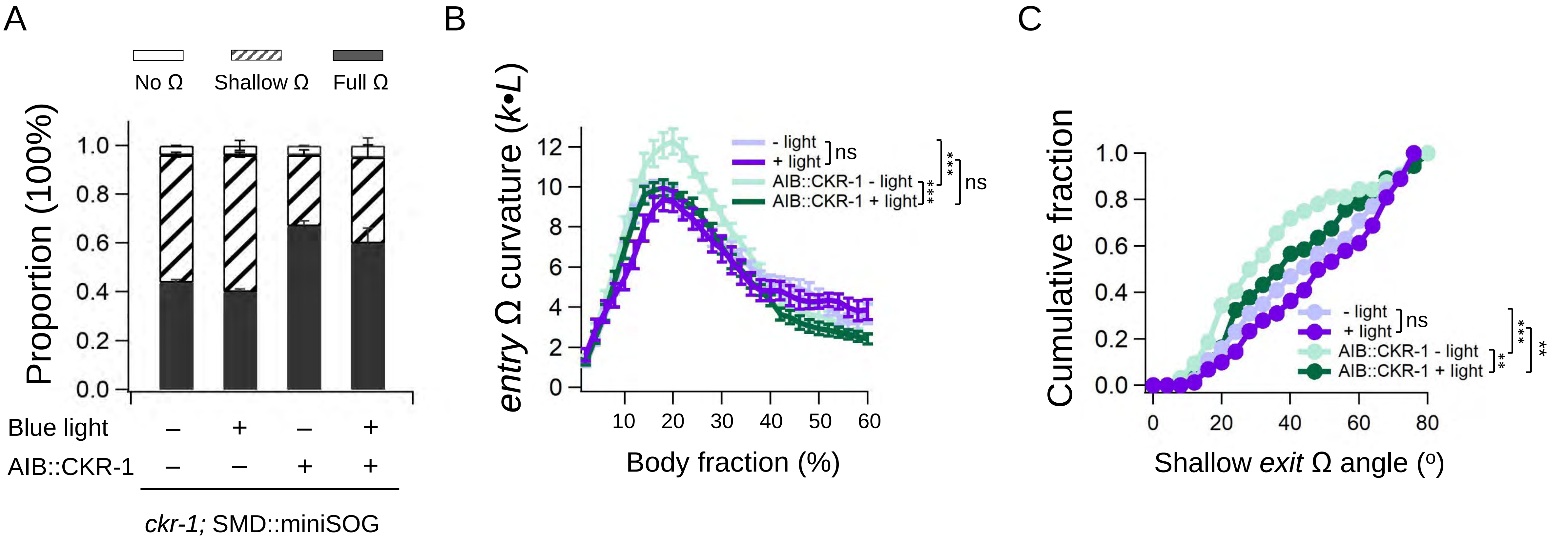
